# PepSpace: An Automated, Physics-Driven Directed Evolution Platform for the De Novo Design of Specific Peptide Binders and Condensate Modulators

**DOI:** 10.64898/2026.09.18.752684

**Authors:** Javier Oller-Iscar, Andrés R. Tejedor, Alejandro Feito, Maria Velasco-Estevez, Alberto Ocaña, Rosana Collepardo-Guevara, Jorge R. Espinosa

## Abstract

Intrinsically disordered proteins, topologically complex protein surfaces, and biomolecular condensates regulate essential cellular processes but remain difficult to drug with conventional small molecules. Here, we present PepSpace, a physics-driven directed-evolution platform coupling residue-resolution coarse-grained molecular dynamics with a multi-objective genetic algorithm. PepSpace designs peptide binders by rewarding target engagement while penalizing peptide self-association and off-target binding. Evaluated alongside deep-learning generative models across multiple targets, PepSpace-designed peptides achieve predicted interaction strengths up to two orders of magnitude greater than machine-learning designs, while reproducing experimentally observed binding hierarchies and revealing distinct binding modes. For PIEZO1, PepSpace designs 25-mer peptides that preferentially engage a defined 30-residue intracellular beam epitope while suppressing interactions with the remainder 2,547-residues of the protein. For CHERP biomolecular condensates, dual-action peptides weaken native intermolecular contacts and destabilize the condensed phase. PepSpace provides a physically grounded framework for de novo peptide design against dynamic protein targets and biomolecular condensates, with explicit control over specificity, solubility, and collective interactions.

## INTRODUCTION

Flexible, non-catalytic regions of the proteome— extended protein–protein interfaces, shallow or topologically complex binding surfaces, and intrinsically disordered proteins/regions (IDP/Rs) that lack a stable folded structure—encompass a large fraction of disease-relevant biology, yet remain among the hardest targets in drug discovery^1^. Conventional small-molecule pharmacology was built around deep, rigid, hydrophobic binding pockets^2^, and that paradigm breaks down when the target is instead a large, solvent-exposed surface or a conformationally plastic region with no pre-organised cavity^3,4^. Monoclonal antibodies can engage such shallow, extended surfaces with high specificity, but their molecular dimensions restrict membrane traversal and access to intracellular targets ^5,6^. Synthetic peptides offer an attractive intermediate modality: conformationally plastic enough to adapt to dynamic, non-planar topologies, large enough to cover extensive contact interfaces, and able to mimic natural regulatory motifs with high affinity. These properties have contributed to the growing use of peptide therapeutics across oncology and other disease areas^7–9^.

Designing peptide binders against dynamic, non-pocket targets is, however, intrinsically multi-objective^10,11^. First, candidate peptides must be evaluated within thermodynamic conformational ensembles rather than against individual static structures, so that target heterogeneity, solvation and entropic contributions to binding can be taken into account^12–14^. Second, on large multi-domain proteins or crowded cellular environments, peptides designed without explicit negative selection can readily accumulate on promiscuous hydrophobic and aromatic patches, resulting in severe off-target cross-reactivity^15–17^.

Biomolecular condensates formed by liquid–liquid phase separation (LLPS) of IDPs, multidomain proteins and nucleic acids are a particularly demanding instance of this problem^18,19^. These assemblies organise spatiotemporal biochemistry within cells^20,21^, from premRNA splicing^22,23^ and transcriptional initiation at super-enhancers^24^ to ribosome biogenesis^25,26^ and the compartmentalisation of DNA-damage repair machinery^27^. Yet the same physical chemistry that enables their formation— weak, multivalent interactions distributed across flexible biomolecules—renders them susceptible to pathological dysregulation^28^. Mutations, post-translational modifications or altered expression can drive aberrant condensation and liquid-to-solid transitions towards cytotoxic, amyloid-like states, as seen for disease-associated variants of the ALS-linked proteins FUS^29^, TDP-43^30,31^ and hnRNPA1^32,33^, which accelerate pathological condensate ageing^14,28,34^. In cancer, dysregulated condensates sustain oncogenic transcriptional programmes, reorganise genome architecture and buffer tumour cells against metabolic and therapeutic stress^28,35,36^, whereas loss of condensation disables tumor suppressors such as UTX and SPOP^37,38^. Selectively correcting these aberrant condensate states—whether by targeted dissolution, stabilization, or restoration—therefore remains a high-priority and largely unmet therapeutic objective^9^. For dissolution, this sharpens the design constraints above: a peptide must engage the very motifs that stabilise the native interaction network while remaining soluble enough to weaken that network rather than be absorbed into it. Furthermore, partitioning into a condensate is not the same as dissolving it.

Deep-learning sequence models such as pepMLM^39^, together with structural generative approaches built on advances in protein structure prediction^40^ such as RFdiffusion^41^, have greatly expanded the scope of *de novo* protein and peptide design. However, these methods do not generally represent solvent effects, ionic screening or conformational fluctuations as explicit components of their design objective^42,43^, which makes highly flexible or disordered targets especially challenging^44,45^. In addition, they do not naturally accommodate competing objectives such as specificity and solubility^46^. Generated candidates consequently require substantial downstream experimental filtering and post-hoc optimisation^47,48^.

Here we introduce PepSpace, an automated, physics-informed discovery platform that couples high-throughput residue-resolution coarse-grained molecular dynamics (MD) simulations^49–52^ with a multi-objective genetic algorithm (GA)^53^ for iterative sequence optimisation. Short MD trajectories serve directly as fitness evaluators, so that peptides are evolved against conformationally dynamic targets while multivalent interactions and sequence-dependent energetics are captured at residue resolution^34,52^. The fitness function in PepSpace rewards target engagement while explicitly penalising peptide self-association and interactions with user-specified off-target regions. We demonstrate this architecture on three distinct design problems. First, we evaluate PepSpace head-to-head against pepMLM^39^ and RFdiffusion^41^ on previously characterised targets, and measure predicted interaction energies up to two orders of magnitude more favorable than those of the generative baselines. Second, we apply it to precision epitope targeting on PIEZO1, a 2,500-residue metastasis-associated mechanosensitive ion channel, designing peptides that selectively engage a designated functional loop while suppressing binding elsewhere on the protein. Finally, we design dual-action peptides against the principal condensation hot spot of the oncogenic splicing factor CHERP. The resulting sequences weaken the intermolecular network driving CHERP phase separation, lower the critical solution temperature and induce condensate dissolution. Together, these results establish PepSpace as a general, physics-informed method to design selective, soluble peptide binders against dynamic and conventionally hard-to-drug targets.

## RESULTS

### PepSpace: A Physics-Driven Platform for Peptide Directed Evolution

PepSpace is an automated, closed-loop directed-evolution framework that couples high-throughput residue-resolution coarse-grained MD simulations ^52^ with a multi-objective genetic algorithm^53^ (Fig. 1). In contrast to sequence- or structure-based generative models that operate primarily on static representations, PepSpace evaluates each candidate across a conformational ensemble generated by molecular dynamics simulations. This makes the platform particularly well suited to flexible, structurally heterogeneous targets, including IDPs, multi-domain proteins, shallow multi-domain surfaces, and other topologically complex, non-planar interfaces that elude conventional structure-based design. The PepSpace discovery pipeline proceeds through five automated stages (Fig. 1).

**FIG. 1.**
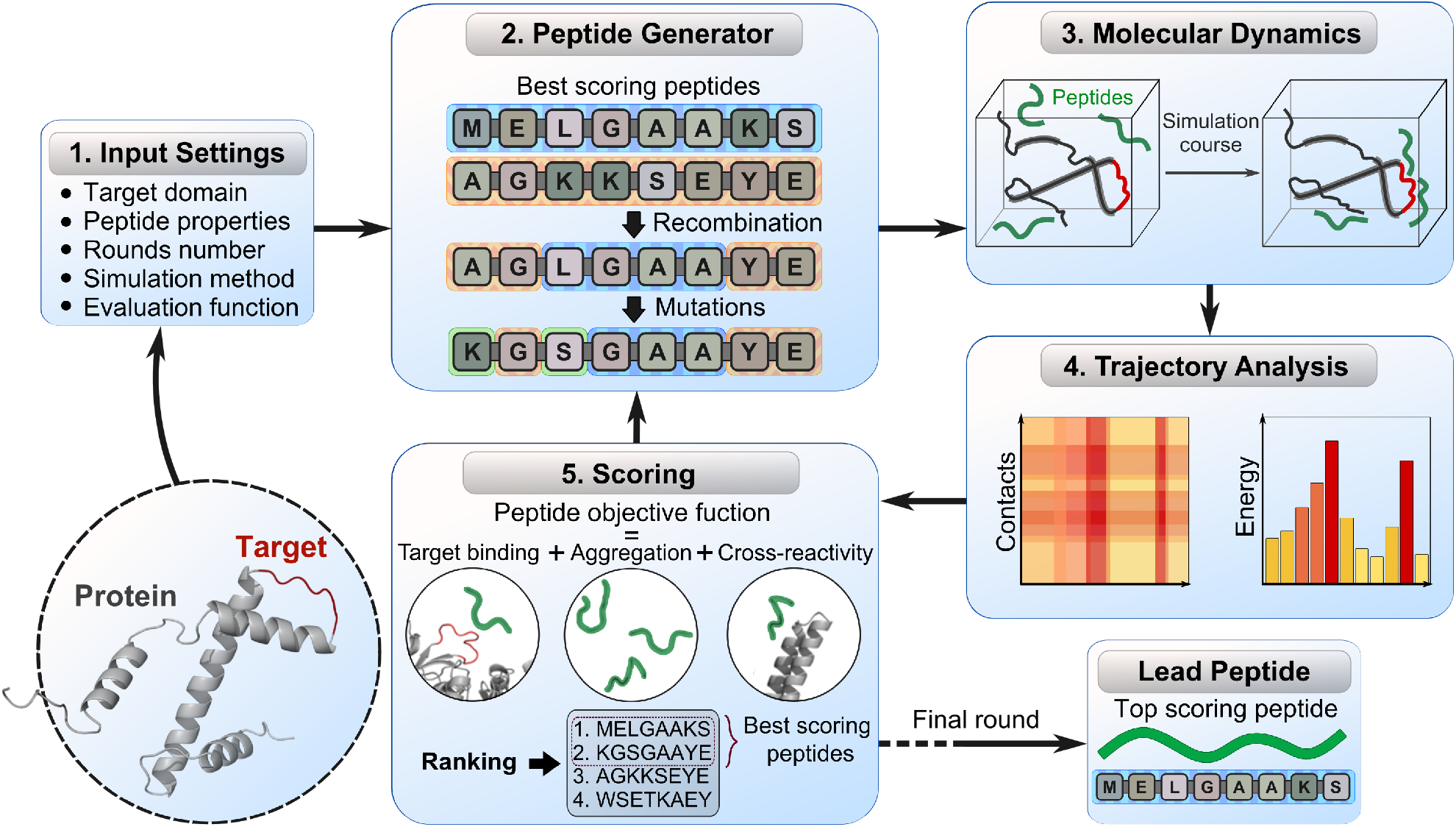
Schematic overview of the automated PepSpace peptide design and discovery platform. Directed evolution operates in a closed iterative cycle: (1) **Target Selection and Input Settings**: Identification of the target protein region (e.g., an intrinsically disordered region, IDR) and definition of configuration parameters, including peptide length, generation size, simulation conditions, and multi-objective Pareto weights. (2) **Peptide Generator**: Construction of the initial sequence population or generation of offspring from top-scoring parents via contact-aware recombination and dynamic mutation operators. (3) **Molecular Simulations**: High-throughput evaluation of candidate peptides within physical simulation boxes containing the target protein under coarse-grained representation. (4) **Analysis**: Quantitative extraction of physical observables, including residue-level intermolecular contact maps and per-residue interaction energies from trajectory ensembles. (5) **Scoring and Ranking**: Multi-objective fitness evaluation using Pareto dominance sorting to identify non-dominated trade-off fronts balancing target binding, homotypic self-aggregation penalties, and off-target cross-reactivity, followed by weighted scalarization within fronts. The top-scoring candidates feed back into the Peptide Generator for subsequent cycles until convergence, yielding the final **Lead Peptide**.

#### Target definition and configuration (Step 1)

The target biomolecule is specified structurally and topographically, including the region or epitope to be engaged and any relevant structural constraints. Additional design parameters specify candidate peptide length, population size per generation, simulation conditions (force field, ionic strength, temperature, and whether simulations are performed with single proteins or biomolecular condensates), and the objective weights that balance target affinity against peptide self-aggregation and cross-reactivity.

#### Peptide generation and genetic operators (Step 2)

The initial population (*t* = 0) is generated under predefined physicochemical and developability constraints intended to restrict the search to therapeutically tractable sequence space. These include modular developability filters—enforcing balanced formal net charge, physiological hydropathy (GRAVY), high aqueous solubility (based on the amount of polar/charged residues), and capping each individual aromatic residue type (Trp, Phe, Tyr) to limit non-specific entropic adhesion and immunogenicity^54,55^. The algorithm additionally allows users to configure structural sequence filters, such as penalizing contiguous tracts of identical residues or hydrophobic amino acids to avoid kinetic aggregation. In subsequent generations (*t >* 0), offspring arise from two genetic operators acting on top-ranked parents: (1) *Contact-aware recombination*, in which crossover breakpoints are guided by residue-level contact and energetic profiles from parent MD short trajectories to preserve beneficial binding motifs; and (2) *Dynamic mutation*, in which point-mutation probabilities are modulated by per-residue target-contact strength so that critical binding residues are conserved while non-interacting positions explore a broad sequence space under the same developability filters.

#### Physics-based MD engine (Step 3)

Each candidate peptide is simulated together with the target protein using the residue-resolution, coarse-grained Mpipi-Recharged force field^52^ in LAMMPS^56^. In contrast to static single-structure or sequence-only generative models, the physical simulation engine samples key determinants of molecular recognition, including conformational flexibility, multivalent interactions, and sequence-dependent energetics. First, target binding objective (*f*_bind_) evaluates peptide–target interaction strength across conformationally heterogeneous trajectories, thereby incorporating effects of structural flexibility, multivalent contacts, and dynamic rearrangements under physiological electrolyte conditions (150 mM NaCl) and defined temperature. Second, homotypic self-aggregation (*f*_agg_) is evaluated directly by simulating multiple peptide copies in the simulation cell, probing whether peptides undergo self-association and phase separation or remain stably dispersed in the implicit aqueous solvent. Third, off-target cross-reactivity (*f*_cross_) is determined by sampling whole-protein surfaces and non-target domains, directly identifying and penalizing non-specific sticking mediated by opportunistic hydrophobic or electrostatic patches (further details on the simulation parameters, system dimensions, and the Mpipi-Recharged potential are provided in Methods and SM Section SII, Equations S1–S6 and Tables S2–S4).

#### Trajectory analysis and energetic decomposition (Step 4)

Trajectory ensembles are processed automatically into distance-based intermolecular contact probability maps and a per-residue energetic decomposition into electrostatic, hydrophobic, and van der Waals contributions (see Methods and SM Section SIII, Equations S7–S10 for calculation details).

#### Multi-objective scoring and lead selection (Step 5)

Candidate sequences are first evaluated using Pareto dominance sorting to identify non-dominated trade-off fronts between competing physical criteria: target binding (*f*_bind_), homotypic self-aggregation (*f*_agg_), and off-target cross-reactivity (*f*_cross_). To rank candidates within each Pareto front and prioritize parent selection, a weighted-sum scalarization is applied:

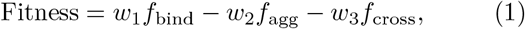

where *f*_bind_ quantifies target interaction energy (contact density and negative interaction potential energy), *f*_agg_ penalizes homotypic peptide-peptide self-association that would otherwise drive aggregation or precipitation^48,57^, and *f*_cross_ penalizes non-specific binding to off-target regions or non-cognate proteins, with *w*_1_, *w*_2_, and *w*_3_ representing objective weighting factors that balance binding efficacy against developability and specificity penalties. Top-ranked candidates across leading Pareto ranks are selected as parents for the next generation, and the cycle repeats until convergence, yielding an optimized lead peptide.

The evolutionary campaigns reported in this work were executed using the residue-resolution Mpipi-Recharged force field^52^, a single-bead-per-residue implicit-solvent model that combines short-range Wang-Frenkel contacts with screened Yukawa electrostatics capturing associative salt-bridge formation. Because the PepSpace directed-evolution algorithm operates as an independent wrapper around the simulation engine, the framework is modular and can be coupled to alternative simulation potentials that provide compatible interaction-energy and contact observables. These include, for instance, alternative coarse-grained models (HPS, CALVADOS, Martini 3) or atomistic explicit-solvent potentials. See Methods and Discussion for complete formulations and transferability.

### Comparative Evaluation of PepSpace Directed Evolution Against Generative Deep-Learning Models

To evaluate the predictive fidelity and design capability of PepSpace, we evaluated it head-to-head against published experimental and computational baselines from state-of-the-art generative models. Specifically, we compare PepSpace against the masked language model *pepMLM* and the structural generative platform *RFdiffusion* (RFD), as reported by Chen et al.^39^, who designed and experimentally validated linear peptide binders—via absorbance assays at 250 nm—for the extracellular domains of two clinically relevant receptors: anti-Müllerian hormone receptor type 2 (AMHR2), an emerging immunotherapeutic target in ovarian cancer^58^, and neural cell adhesion molecule 1 (NCAM1/CD56), a validated antibody-drug-conjugate target in small-cell lung cancer and other CD56-positive malignancies^59^.

We first conducted a calibration and validation of the simulation framework by examining how published machine-learning-derived peptides bind in explicit thermodynamic ensembles. To benchmark these designs systematically, we simulated candidate peptides reported by Chen et al.^39^ across both targets (Fig. 2a,b, SM Fig. S3, and SM Table S5), comparing the simulated binding behavior of the best and worst experimental performers with the Mpipi-Recharged force field. For the compact extracellular domain of AMHR2 (Fig. 2a), the simulated interaction energy per residue reflects the experimental contrast observed between the top *pepMLM* peptide (*pepMLM-1, AMHR2-pepMLM-03, A*_250_ *≈* 0.52) and the weakly active *RFdiffusion* design (*RFD-1, AMHR2-RFD-01, A*_250_ *≈* 0.06). For NCAM1 (Fig. 2b), the simulations similarly capture baseline target interactions across the candidate designs. Having established that the Mpipi-Recharged model—already extensively benchmarked and validated across more than 100 protein systems in independent studies^52,60^—provides a physically sound and consistent landscape for peptide evaluation, we then deploy this force field within PepSpace to guide sequence optimization. Crucially, under this physical objective, PepSpace explores regions of sequence space that generative models do not reach, uncovering alternative interaction modes and evolving candidate peptides that bind substantially stronger to both targets than any of the generative benchmarks.

**FIG. 2.**
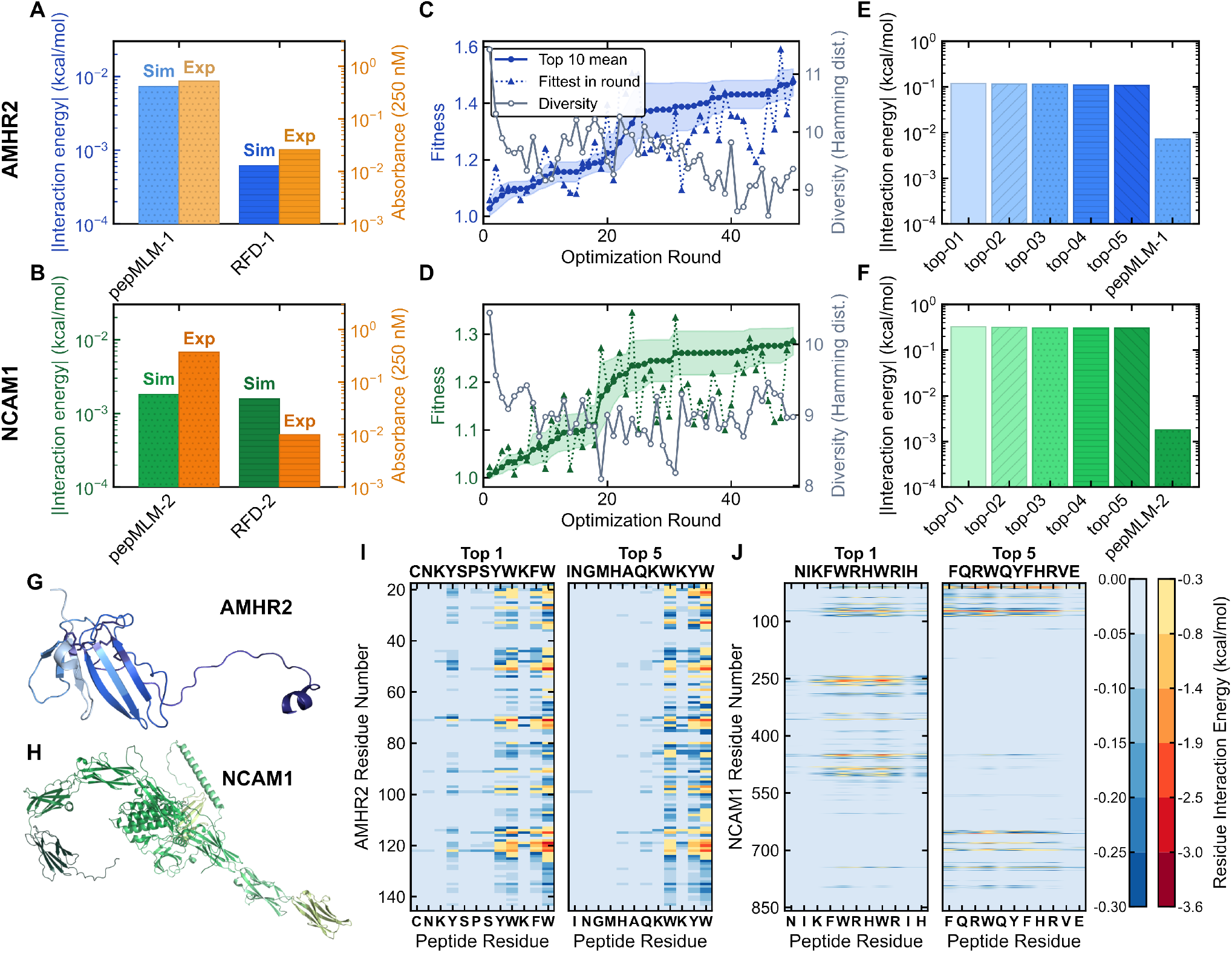
Benchmarking and comparative evaluation of PepSpace against *pepMLM* and *RFdiffusion* across AMHR2 and NCAM1. **a**,**b**, Experimental benchmarking of coarse-grained molecular dynamics interaction energies normalized per peptide residue (blue/green bars, left axis, in kcal/mol per amino acid) against experimental absorbance measurements at 250 nm (*A*250, orange bars, right axis) from Chen et al.^39^ for the top *pepMLM* and *RFdiffusion* (RFD) designs targeting the extracellular domains of AMHR2 (**a**) and NCAM1 (**b**). **c**,**d**, Directed evolutionary trajectory over 50 rounds (15 candidate peptides evaluated per round) showing the mean composite fitness of the top-10 candidates (solid line, shaded band represents *±* 1 standard deviation), the round best candidate (triangles), and population sequence diversity measured by mean pairwise Hamming distance (grey line) for AMHR2 (**c**) and NCAM1 (**d**). **e**,**f**, Comparison of absolute interaction energies per peptide residue (|Interaction energy|, kcal/mol per amino acid, log scale) between the top-5 evolved peptides from PepSpace and the best experimental *pepMLM* binder for AMHR2 (**e**) and NCAM1 (**f**). **g**,**h**, Structural models of the extracellular target domains of AMHR2 (**g**) and NCAM1 (**h**). **i**,**j**, Two-dimensional residue-level contact interaction energy matrices (*Eij*, kcal/mol) resolving pairwise contacts between target extracellular residues (y-axis) and peptide amino acid positions (x-axis) for the *top-1* and *top-5* evolved candidates binding AMHR2 (**i**) and NCAM1 (**j**).

Because binding free energies are intrinsically state-dependent functions of temperature, ionic screening, and conformational ensemble heterogeneity, physics-based molecular dynamics simulations allow these variables to be included explicitly during candidate evaluation. PepSpace therefore adds a distinct physical selection layer to generative sequence and structure design, evaluating conformational fluctuations, solution conditions, self-association, and off-target interactions directly rather than relying on static structural representations or learned sequence priors alone. For instance, in dynamic or membrane-adjacent targets such as the mechanosensitive channel PIEZO1, binding is intimately governed by local electrostatic conditions and conformational flexibility. Applying structural diffusion models such as RFdiffusion or RFD-2^41^ to such systems presents major challenges. These architectures are primarily trained on folded protein structures, scale unfavourably with very large, multi-thousand-residue targets, and are difficult to apply to intrinsically disordered loops that lack a stable single-conformation coordinate frame. They also do not explicitly capture the thermodynamic effects of solvation, ionic conditions, or negative selection against the surrounding protein surface during binder design.

Having validated the simulation engine, we now deploy PepSpace’s full directed-evolution cycle over 50 rounds (15 candidate peptides per round), which produced a progressive increase in mean population fitness (Fig. 2c,d and SM Fig. S1) for both AMHR2 and NCAM1 sequences. The dynamic mutation operator preserved broad sequence diversity throughout optimization—maintaining a mean pairwise Hamming distance of *∼* 8.5–9.5—preventing premature convergence and enabling thorough exploration of the sequence space. Comparing the equilibrium binding energetics of the top-5 evolved peptides against the *pepMLM* lead candidates reveals substantially stronger predicted peptide–target interaction energies (SM Fig. S2 and SM Table S6). On AMHR2 (Fig. 2e), the top-5 PepSpace candidates reached interaction energies of *∼ −* 0.11 kcal/mol per residue, exceeding the designed *pepMLM-1* baseline (*≈ −* 0.007 kcal/mol per residue) by more than an order of magnitude. On NCAM1 (Fig. 2f), evolved peptides attained interaction energies below −0.30 kcal/mol per residue, compared to −0.002 kcal/mol per residue for *pepMLM-2* —an enhancement exceeding two orders of magnitude in simulated binding strength.

Residue-level contact interaction matrices (Fig. 2i,j) reveal how the evolutionary engine autonomously adapts its search strategy to contrasting target topologies. On the compact, single-domain fold of AMHR2 (*<* 150 residues, Fig. 2g), the *top-1* (*CNKYSPSYWKFW*) and *top-5* (*INGMHAQKWKYW*) evolved peptides converged on a shared C-terminal aromatic motif (−WKFW and -WKYW) that anchors cooperatively into the central *β*-sheet and adjacent flexible loops (residues 45–55, 70– 80, and 115–125 with pairwise interaction energies *< −* 3.0 kcal/mol; Fig. 2i). On the expansive, multi-domain architecture of NCAM1 (*>* 700 residues spanning multiple immunoglobulin-like domains, Fig. 2h), by contrast, PepSpace bypassed localized traps to discover distinct binding modes: the *top-1* peptide (*NIKFWRHWRIH*) concentrated its interactions on the Ig2/Ig3 hinge via an arginine-tryptophan triad (FWRHWR), whereas the *top-5* peptide (*FQRWQYFHRVE*) engaged multiple domains simultaneously, bridging Ig1 and the membrane-proximal stalk (Fig. 2j). This divergence shows that whole-surface evolutionary search can uncover topologically distinct, comparably favourable binding solutions rather than converging on a single dominant interaction pattern (Fig. 2f). By evaluating candidate sequences directly across dynamic target surfaces, PepSpace extends the objective space of current sequence- and structure-generative models and allows alternative binding modes to emerge from the underlying physical interaction landscape.

A key question for any *de novo* peptide design platform is whether increasing target interaction strength comes at the cost of specificity: optimizing peptides for target interaction strength alone, without explicitly accounting for off-target interactions, can produce substantial cross-reactivity. To examine this issue, we evaluate binding of our top evolved leads, as well as the benchmark *pepMLM* designs, against both AMHR2 and NCAM1 (Fig. 3a,b and SM Table S7). Our designed peptides achieve strong on-target interaction strength but also show notable cross-reactivity. Peptides evolved against NCAM1 bound off-target AMHR2 nearly as effectively as the cognate AMHR2-designed peptide, and the converse held true on NCAM1. The baseline *pepMLM* lead peptides, by comparison, showed overall weaker simulated interactions but a markedly more pronounced lack of specificity: *pepMLM-1*, designed for AMHR2, bound offtarget NCAM1 more favorably than its intended target— and even outperformed *pepMLM-2*, the peptide actually designed for NCAM1, at binding NCAM1 itself. Such off-target interactions would represent a potential translational liability if retained experimentally, as non-specific pharmacology is a leading cause of safety-related attrition in drug development^61^.

**FIG. 3.**
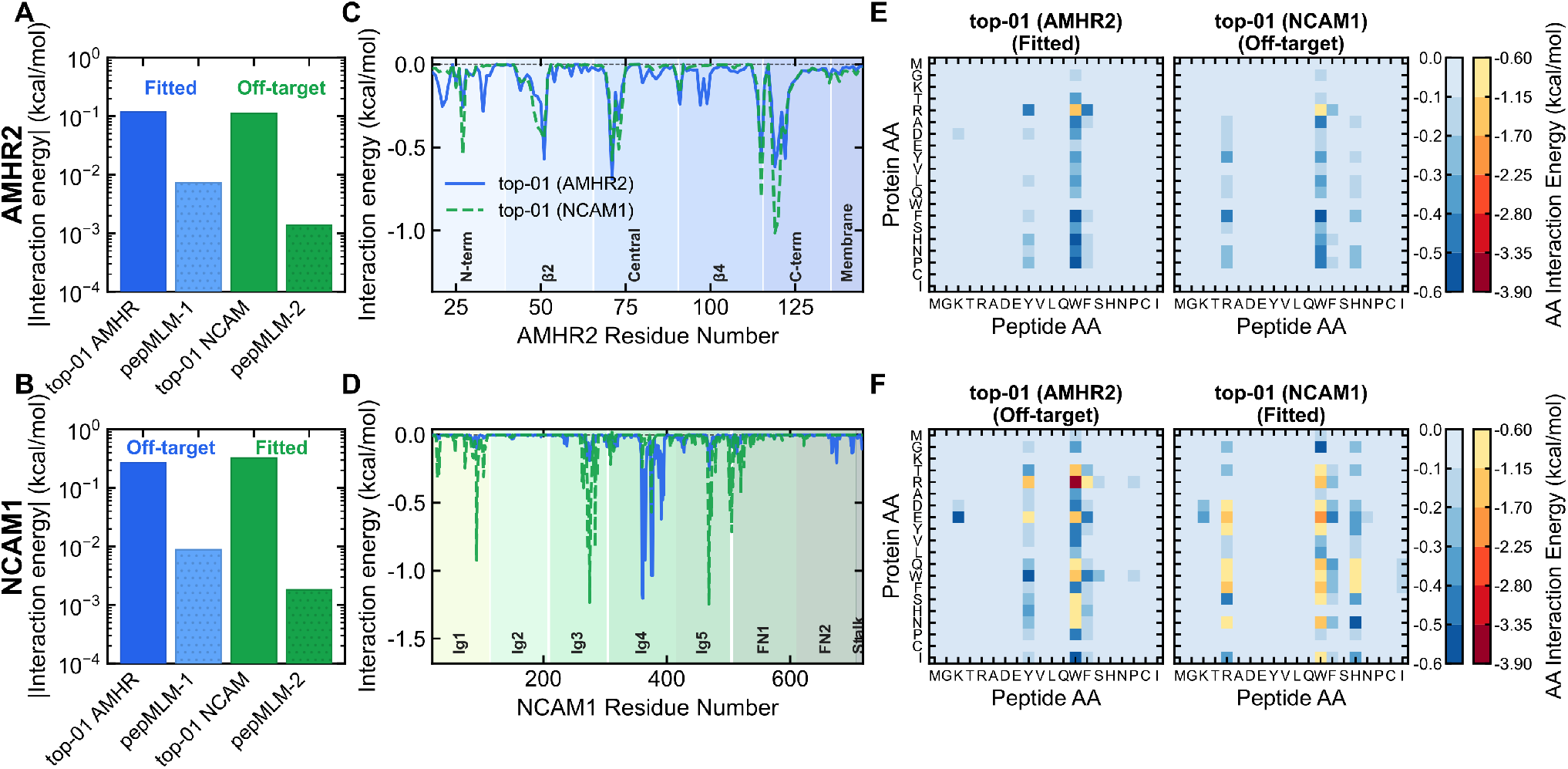
Cross-reactivity and binding specificity analysis between AMHR2 and NCAM1 designed peptides. **a**,**b**, Comparison of absolute interaction energies normalized per peptide residue (|Interaction energy|, kcal/mol per amino acid, log scale) for on-target (fitted) versus off-target cross-reactive binding across AMHR2 (**a**) and NCAM1 (**b**), evaluated for both our evolved *top-1* peptides and the *pepMLM* benchmark candidates. **c**,**d**, Per-residue interaction energy profiles along the primary sequence of the target receptors (*E*(*i*), kcal/mol per target residue) for AMHR2 (**c**) and NCAM1 (**d**) engaging *top-01 (AMHR2)* (blue solid curve) and *top-01 (NCAM1)* (green dashed curve), with protein domain boundaries indicated. **e**,**f**, Two-dimensional amino acid interaction energy contact matrices resolving pairwise contacts between target protein residue types (y-axis) and peptide amino acid types (x-axis) for on-target and off-target complexes on AMHR2 (**e**) and NCAM1 (**f**).

To understand at molecular level how this cross-reactivity arises across such different receptor architectures, we examine the per-residue interaction profile along the sequence of each target (Fig. 3c,d). On AMHR2 (Fig. 3c), the cognate and off-target peptides produce nearly superimposable interaction profiles, both converging on the same structural features—the *β*2 loop, the central *β*-sheet, and the flexible C-terminal segment—a similarity explained by the compact size of the AMHR2 extracellular domain, whose limited surface area funnels distinct peptides toward the same accessible binding region. On the expansive, multi-domain architecture of NCAM1 (Fig. 3d and SM Table S8), by contrast, our two peptides partition into entirely distinct structural domains: the off-target AMHR2 peptide binds almost exclusively to the Ig4 domain, whereas the cognate NCAM1 peptide leaves Ig4 largely uncontacted and instead distributes its binding across the Ig1, Ig3, and Ig5 domains.

The underlying chemistry of these profiles is resolved by the two-dimensional amino-acid contact matrices (Fig. 3e,f). On AMHR2 (Fig. 3e), both the cognate and offtarget peptides rely on a shared mechanism, in which peptide tryptophan residues make the dominant contribution through strong interactions with positively charged and aromatic residues on the target. On NCAM1 (Fig. 3f), the interaction chemistry instead diverges: tryptophan again contributes strongly in both peptides, but the off-target AMHR2 peptide additionally engages through tyrosine, whereas the cognate NCAM1 peptide instead recruits arginine and polar histidine residues to build a distinct electrostatic contact network. Together, these results show that binder optimization without counter-selection naturally exploits whatever hydrophobic and aromatic surface features are accessible, regardless of target identity, underscoring why explicit negative design against off-target proteins is essential to achieve greater target selectivity rather than interaction strength alone, as we will show in next sections.

Importantly, this comparative evaluation reflects the fundamentally distinct paradigms of the two methodologies, each presenting distinct advantages. Deep-learning generative platforms feature exceptional throughput: sequence-based masked language models (such as *pepMLM*) and structural diffusion frameworks (such as *RFdiffusion*) evaluate sequence logits or structural backbones in seconds to minutes on GPU hardware, enabling rapid, unconstrained zero-shot sequence exploration across uncharacterized sequence spaces. In contrast, PepSpace trades instantaneous sequence proposal for explicit, physically grounded thermodynamic sampling. Because PepSpace evaluates candidate populations directly across microsecond-scale conformational ensembles and extracts multi-objective physical fitness through molecular dynamics, it is inherently several orders of magnitude more computationally demanding than purely predictive or static machine-learning inference (see Methods and SM Section SIV for complete simulation protocols, parallelization benchmarks, and wall-clock runtimes). In return for this computational investment, however, PepSpace provides direct access to physical observables that are not usually included explicitly in sequence- or structure-generative models: direct sampling of conformational fluctuations under physiological electrolyte conditions, residue-level energetic partitioning, and the ability to enforce rigorous multi-objective counter-selection against self-aggregation and off-target cross-reactivity. Rather than competing paradigms, these approaches represent complementary discovery strategies, where rapid generative sequence proposal could seamlessly feed into physics-driven Pareto refinement.

### Site-Specific Peptide Targeting within the PIEZO1 *>* 2500-residue Protein

Having established peptide-protein whole-surface optimization, we next challenge PepSpace to achieve precision, site-specific targeting within a large, multi-domain macro-molecular protein under active therapeutic scrutiny: the mechanosensitive ion channel PIEZO1 (*>* 2500 residues per monomer, Fig. 4a)^62–64^. This homotrimeric channel couples shear-stress-activated calcium conduction to vascular remodeling^65^ and to the mechanotransduction program that drives tumor cell migration, invasion, and metastatic extravasation^66^, making it an actively pursued oncology and cardiovascular target. Structural studies have established that the intracellular helical beam acts as a rigid mechanical lever, coupling blade deflection to pore gating^64^. Existing chemical modulators of PIEZO1, such as the small-molecule agonist Yoda1, act directly at the blade–pore interface^67^. We instead will target a solvent-exposed, conformationally dynamic 30-residue loop within the beam (residues 1373–1402) as an orthogonal, non-pore-blocking target epitope (Fig. 4b), offering a route to modulate channel mechanosensitivity without directly occluding in a non-specific manner the conduction pathway. Designing a peptide binder for this region poses a severe biophysical challenge: the candidate must selectively recognize a dynamic 30-residue segment while ignoring the remaining 2500 residues of competing hydrophobic transmembrane helices, blade repeats, and intracellular loops.

**FIG. 4.**
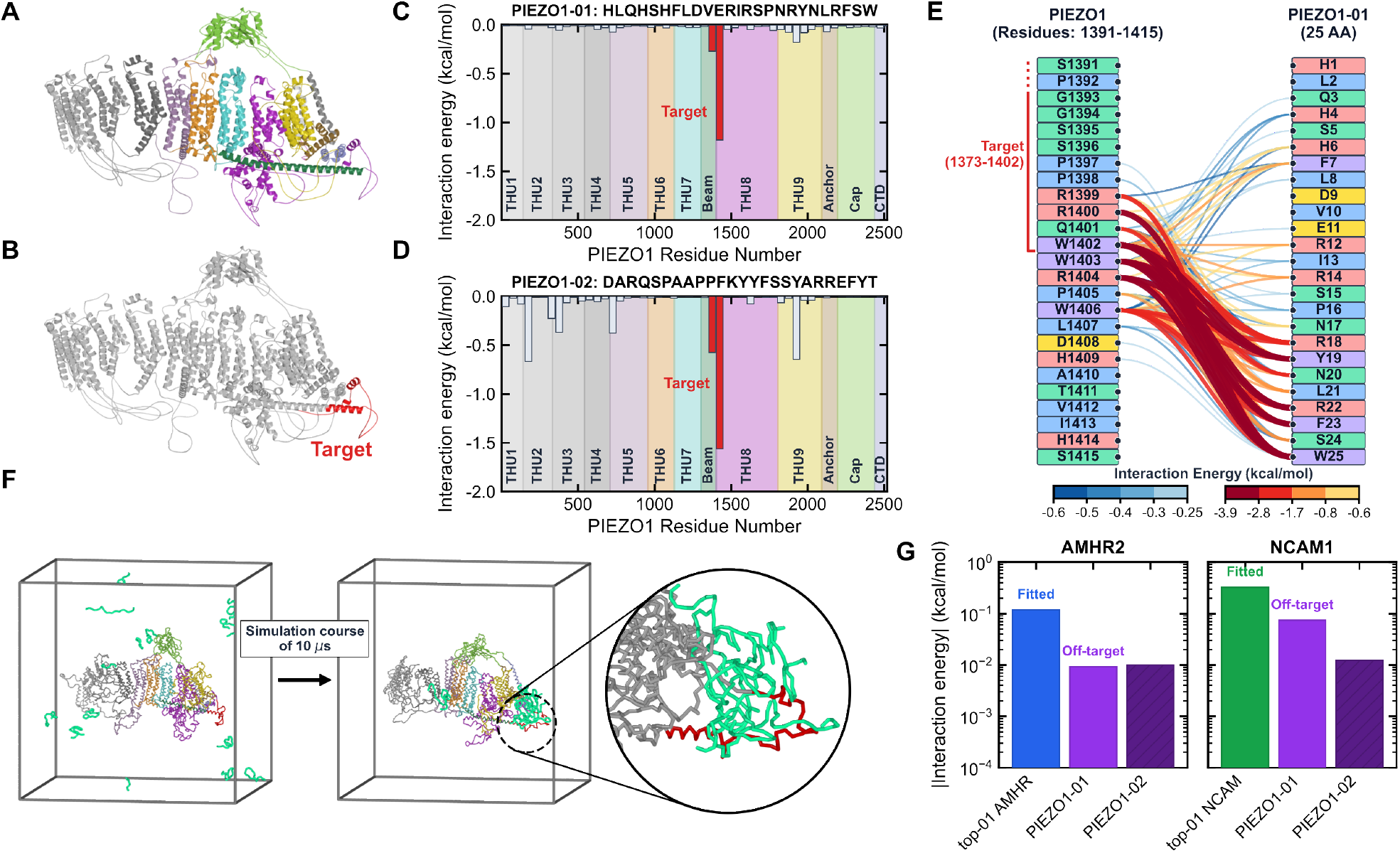
Targeted de novo design of high-affinity, specific peptide binders for the PIEZO1 beam domain. **a**, Structural architecture of the full-length human PIEZO1 monomer (2547 residues) colored by structural domains (THU1–THU9 blades, Beam, Anchor, Cap, and CTD)^63,64^. **b**, Structural localization of the designated 30-residue target epitope (residues 1373–1402, highlighted in red) within the intracellular helical beam domain. **c**,**d**, Whole-protein per-residue interaction energy profiles along the complete 2547-residue sequence of the PIEZO1 monomer for the top two evolved 25-mer peptides, *PIEZO1-01* (*HLQHSHFLDVERIRSPNRYNLRFSW*) (**c**) and *PIEZO1-02* (*DARQSPAAPPFKYYFSSYARREFYT*) (**d**), evaluated in 50-residue sliding windows (Interaction energy, kcal/mol per window), confirming exclusive, site-specific binding to the designated target epitope with near-zero background off-target interactions across the remaining channel body. **e**, Bipartite residue-to-residue interaction network diagram resolving pairwise energetic couplings between PIEZO1 beam residues (1391–1415, left nodes, with the designated target segment 1391–1402 demarcated by the red bracket) and individual amino acid positions of the lead peptide *PIEZO1-01* (H1–W25, right nodes); edge colors and line thicknesses reflect the pairwise interaction energy (*Eij*, kcal/mol) with node badges colored by amino acid physicochemical class. **f**, Representative simulation trajectory snapshots showing initial random placement of candidate peptides in the bulk solvent box (*t* = 0, left) and final specific assembly and stable anchoring onto the intracellular beam target zone after equilibration (*t* = 10 *µ*s, right). **g**, Quantitative off-target cross-reactivity comparison showing absolute interaction energies normalized per peptide residue (|Interaction energy|, kcal/mol per amino acid, log scale) of the PIEZO1-designed peptides against non-cognate receptors AMHR2 and NCAM1 relative to their cognate fitted binders.

To this purpose, we now execute PepSpace with negative-selection weights (*w*_3_ *>* 0; see Methods and SM Section SI for further details on multi-objective weighting and constraints) penalizing non-target interactions across the entire PIEZO1 monomer. The resulting 25-mer peptide leads, *PIEZO1-01* (*HLQHSHFLDVERIRSPN-RYNLRFSW*) and *PIEZO1-02* (*DARQSPAAPPFKYYF-SSYARREFYT*), achieve strong spatial specificity on the target protein domain, Fig. 4c and Fig. 4d, respectively. Whole-protein per-residue interaction energy profiles, evaluated in 50-residue sliding windows, show preferential localization to the designated target epitope, with peak interaction energies of −1.25 and −1.60kcal/mol per window (Fig.4c,d and SM Table S9) against a background of near-zero off-target binding reactivity across the vast remaining channel surface.

Residue-level contact-network analysis (Fig. 4e) identifies three complementary features associated with the high target selectivity of *PIEZO1-01*. First, the C-terminal motif Y19-N20-L21-R22-F23-S24-W25 forms strong aromatic and cation-*π* contacts with R1399, R1400, Q1401, and W1402, as well as the adjacent beam residues W1403, R1404, and W1406. Pairwise interaction energies reach −3.8 to −4.5kcal/mol, with particularly strong contributions from W1402/W1403–R22 and R1400/R1404–W25. Second, internal arginine residues R12, R14, and R18 interact with the tryptophan-rich beam cluster W1402, W1403, and W1406, with pairwise energies of −0.6 to −2.7kcal/mol. Third, flexible proline and polar residues, including P16, N17, and S24, support conformational adaptation of the peptide along the helical beam surface, together with additional N-terminal contacts involving H4, H6, and F7. Collectively, these interactions produce a distributed recognition pattern combining aromatic anchoring, electrostatic coordination, and backbone flexibility. This type of multimodal binding is difficult to capture through optimization against a single static target conformation. Consistent with this interaction pattern, peptides initialized randomly in bulk solvent diffuse to the target region and remain preferentially localized at the beam epitope over the simulation trajectory (*t* = 10*µ*s; Fig.4f), with minimal non-specific interaction elsewhere on the channel surface.

Beyond intra-molecular precision, a central question for any targeted therapeutic candidate is whether high on-target interaction strength incurs off-target polyspecificity across the proteome. To address this, we evaluated the cross-reactivity of *PIEZO1-01* and *PIEZO1-02* peptides against the two previously studied proteins: AMHR2 and NCAM1 (Fig. 4g and SM Table S10). Against AMHR2 (Fig. 4g, left), where the cognate fitted peptide (*top-01 AMHR*) achieves an average interaction energy of *∼* 0.12 kcal/mol per residue, the off-target interaction energies of *PIEZO1-01* and *PIEZO1-02* drop to only *∼* 9.0 *×* 10^−3^ and *∼* 1.0 *×* 10^−2^ kcal/mol per residue, respectively—a decrease of more than an order of magnitude. Similarly, against NCAM1 (Fig. 4g, right), where the *top-01 NCAM* peptide achieves an average interaction energy of *∼* 0.33 kcal/mol per residue, off-target binding of *PIEZO1-01* is suppressed to *∼* 7.5 *×* 10^−2^ kcal/mol per residue, while *PIEZO1-02* shows an even more pronounced reduction, to *∼* 1.2 *×* 10^−2^ kcal/mol per residue— a suppression of nearly 1.5 orders of magnitude.

These cross-reactivity profiles stand in sharp contrast to the unconstrained evolutionary designs of Fig. 3a,b, where binders lacking negative selection readily engaged foreign receptors with affinities matching or exceeding those of their cognate leads. Without explicit counter-selection, sequence evolution habitually exploits generic hydrophobic and aromatic patches that adhere opportunistically to any available macromolecular surface. It is important to note, however, that AMHR2 and NCAM1 were not themselves included as negative-selection targets during the evolution of *PIEZO1-01* and *PIEZO1-02*. The only counter-selection pressure applied was against off-target regions within the PIEZO1 monomer itself (Fig. 4c,d). Hence, the low cross-reactivity we observe against these two entirely unrelated folds is an emergent property of specifying a narrow, chemically distinctive target epitope, rather than a directly engineered outcome against these specific proteins. This observation has two implications. First, precision epitope targeting confers a degree of proteome-wide specificity as an intrinsic byproduct, without requiring every possible off-target protein to be enumerated during design. Second, this emergent specificity nonetheless leaves room for further improvement: explicitly incorporating AMHR2, NCAM1, or other candidate off-target proteins into the negative-selection set would be expected to suppress residual cross-reactivity even further, and represents a natural refinement for future PepSpace campaigns targeting clinical candidates. Taken together, these results establish PepSpace’s multi-objective fitness formulation as an effective computational strategy for engineering targeted, clinically relevant peptide modulators against complex, multi-domain protein sequences.

### Engineered Dual-Action Peptides for CHERP-specific Binding and Condensate Disruption

We next use PepSpace to address one of the most demanding challenges in macromolecular therapeutics: the targeted dissolution of a biomolecular condensate via engagement of IDRs. We focused on the oncogenic splicing factor CHERP (916 residues), whose phase separation has been associated with colorectal, esophageal, and gastric carcinomas and with pro-survival alternative splicing programs, consistent with a broader mechanistic link between splicing dysregulation and cancer biology^68,69^. Direct Co-existence simulations of full-length CHERP mapped the intermolecular interaction network that stabilizes the condensed phase (Fig. 5a). Residue-level energetic decomposition revealed a well-defined condensation hotspot within the intrinsically disordered region, spanning residues 476– 576. This segment is enriched in positively charged and aromatic residues and contributes more than 25% of the total intermolecular interaction energy of the full 916-residue protein, identifying it as a dominant driver of CHERP condensation.

**FIG. 5.**
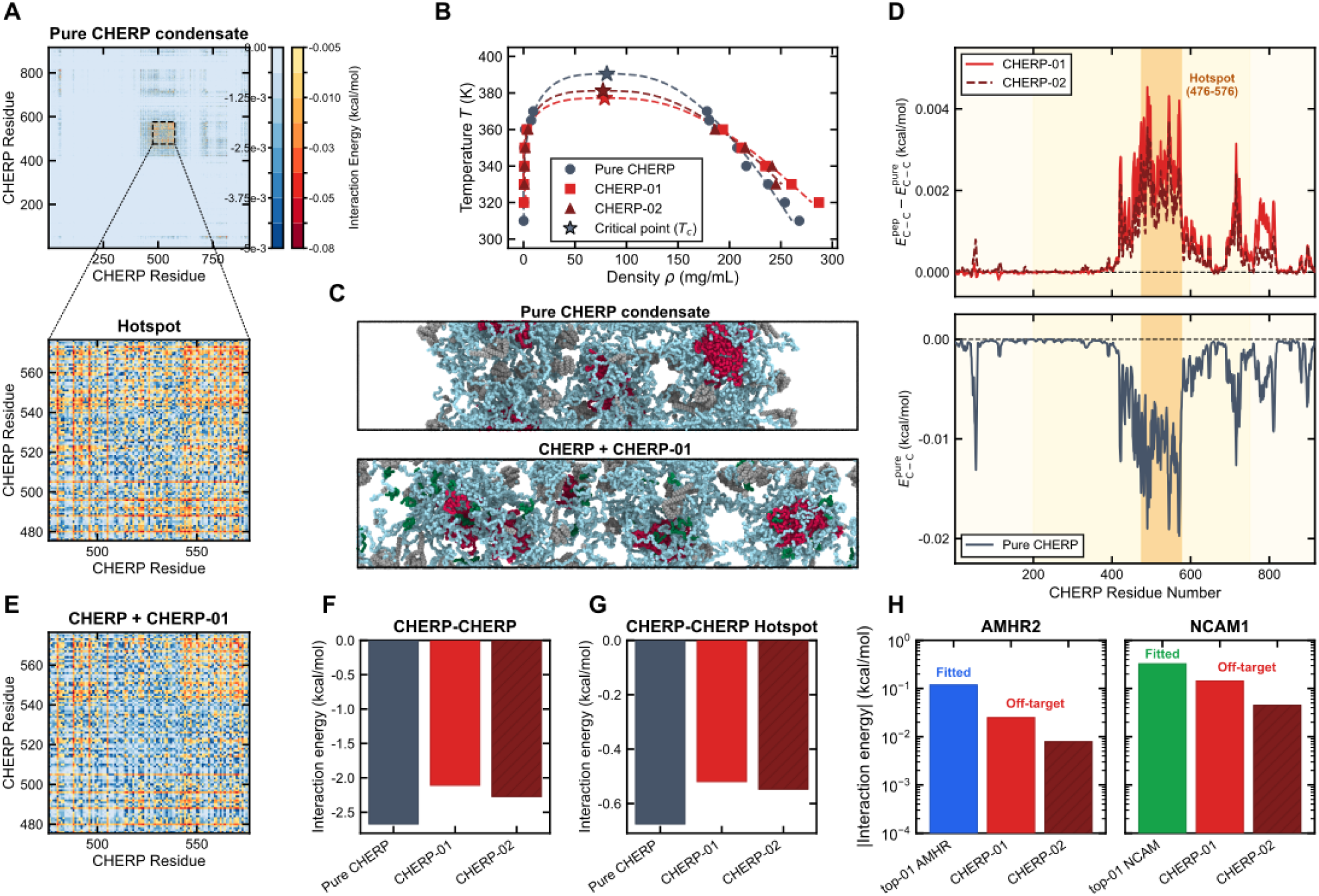
Targeted disruption of CHERP liquid-liquid phase separation via dual-action designed peptides. **a**, Residue-level contact interaction energy map of full-length homotypic CHERP-CHERP interactions at 370 K in the pure condensate (no peptide), revealing a dominant intermolecular interaction hot spot localized to residues 476–576 (inset zoom); dual colorbars show weak (0.00 to −5.0 *×* 10^−3^ kcal/mol, blue) and strong (−0.005 to −0.08 kcal/mol, red) interaction regimes. **b**, Temperature-density coexistence phase diagrams for the 25-molecule CHERP condensate system in the absence of peptide (black circles) and upon addition of 50 molecules of *CHERP-01* (red squares) or *CHERP-02* (brown triangles), highlighting the downward shift in critical temperature (*Tc*, open stars) calculated via direct coexistence simulations^60,70^ fitted with 3D Ising critical scaling and rectilinear diameters. **c**, Direct coexistence slab simulation snapshots at 370 K illustrating the intact multi-core dense CHERP condensate without peptide (top) versus pronounced core disruption and chain dispersion into the dilute phase upon addition of *CHERP-01* (bottom; peptides shown in green, CHERP chains in grey/red/yellow). **d**, Per-residue interaction energy profiles along the 916-residue CHERP sequence at 370 K: top panel displays the differential homotypic interaction energy 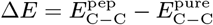, in kcal/mol per residue per pair), where positive values quantify targeted energetic weakening localized to the 476–576 hot spot; bottom panel displays the absolute per-residue homotypic contact energy in the pure condensate (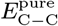, in kcal/mol per residue per pair, smoothed with a 5-residue moving average). **e**, Zoomed contact energy matrix of CHERP-CHERP interactions in the presence of *CHERP-01* within the 476–576 hot spot, illustrating attenuated contact density and an increased fraction of weak blue contacts. **f**,**g**, Mean homotypic intermolecular interaction energy per CHERP–CHERP pair (in kcal/mol per pair across the 300 intermolecular pairs) integrated across the entire 916-residue sequence (**f**) and restricted strictly to contacts within the 476–576 hot spot (**g**) for pure CHERP, *CHERP-01*, and *CHERP-02* systems. **h**, Off-target cross-reactivity comparison confirming low non-specific binding of CHERP peptides against non-cognate receptors AMHR2 and NCAM1 (normalized per peptide residue, log scale).

Importantly, condensate dissolution imposes a dual design requirement that differs from conventional interaction-driven peptide optimization. A peptide optimized solely for strong target binding may co-partition into the dense phase and introduce additional multivalent interactions, thereby stabilizing the condensate or, in some cases, promoting liquid-to-solid transitions^14^. Effective dissolution therefore requires strong peptide– target interactions (*f*_bind_ in Eq.1) to compete with native CHERP–CHERP contacts, together with minimal peptide–peptide self-association (*f*_agg_) and high aqueous solubility. Such peptides can engage and cap interaction hotspots without introducing new cohesive crosslinks into the condensate network. However, low peptide self-association alone does not guarantee condensate disruption. Weakly self-associating peptides can instead accumulate at the condensate interface and behave as molecular surfactants, lowering surface tension without substantially disrupting the internal interaction network^71–73^. PepSpace explicitly balances these competing physical requirements through its multi-objective fitness function. For CHERP, PepSpace identified two lead sequences: *CHERP-01* (30-mer, *GMRMDLQHCLENL-RFPRKTWRHSWKHFKAY*) and *CHERP-02* (25-mer, *YCHLTHSWDHRTWETCHVLFIHYSI*).

We use Direct Coexistence simulations across a wide range of temperatures (300–400 K) to compute the phase diagram of CHERP in both presence and absence of the two generated peptides (Fig. 5b and SM Table S11; see Methods and SM Section SIII, Equations S11–S14 for methodological details^60,70^). Our simulations show a strong destabilization of CHERP condensates by the PepSpace-designed peptides. In the absence of peptides, pure CHERP exhibits a critical solution temperature of *T*_*c*_ = 390.5 K (with critical density *ρ*_*c*_ = 80.9 mg/mL) in our model. The addition of *CHERP-02* decreases the critical solution temperature by Δ*T*_*c*_ = −9.3 K (to *T*_*c*_ = 381.2 K), while *CHERP-01* drives a substantially larger depression of Δ*T*_*c*_ = −13.3 K (down to *T*_*c*_ = 377.2 K). Direct Coexistence simulation snapshots below the critical solution temperature of pure CHERP condensates at *T* = 370 K (corresponding to *T/T*_*c*_ *≈* 0.95 for pure CHERP, but shifted to *T/T*_*c*_ *≈* 0.98 near the critical boundary upon peptide addition. Fig. 5c) visually display such inhibition: *CHERP-01* (dark green chains) intercalates into the condensed phase, breaking CHERP–CHERP multivalent intermolecular contacts and dispersing CHERP proteins into a homogeneous diluted phase (bottom panel).

Residue-level contact profiling (Fig.5d,e) shows that the phase-boundary shift is associated with targeted weakening of the condensation core rather than diffuse disruption across the full protein. The reduction in native CHERP– CHERP contacts 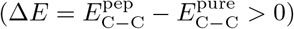 is strongly localized to the 476–576 hot spot, where differential weakening reaches +0.0045 kcal/mol per residue per pair. At the whole-protein level, *CHERP-01* and *CHERP-02* reduce the mean homotypic CHERP interaction energy per intermolecular pair from −2.67 kcal/mol to −2.11 and −2.27 kcal/mol, respectively (Fig.5f). This corresponds to an energetic attenuation of Δ*E ≈* 0.40–0.56 kcal/mol per pair, or approximately 15–21% of the total intermolecular interaction energy.

In multivalent IDR phase separation, the phase boundary reflects a fine balance between chain conformational entropy and collective intermolecular interactions. Changes of only a few tenths of a kcal/mol, for example through single post-translational modifications or disease-associated mutations, have been shown experimentally and computationally to shift saturation concentrations substantially or abolish liquid–liquid phase separation^49,74–76^. Here, the targeted attenuation of up to *∼* 0.56 kcal/mol per pair is accompanied by a decrease in the critical solution temperature of up to 13.3 K (Fig.5b), directly linking disruption of the interaction network to a measurable shift in the phase boundary. Within the 101-residue hot spot alone, homotypic interactions decrease from −0.67 to −0.52 kcal/mol per pair (Fig.5g and SM Table S12). Summing the per-residue interaction energies across the 101 hot-spot residues yields precisely this −0.67 kcal/mol per pair (with individual residue energies in Fig. 5d peaking at −0.020 kcal/mol per pair at residue 571). Despite this strong effect on CHERP condensation, both peptides retain lower interaction with the non-cognate proteins AMHR2 and NCAM1 (Fig.5h and SM Table S13). As in the PIEZO1 design, negative selection was applied only against off-target regions within the CHERP sequence and not directly against AMHR2 or NCAM1. The reduced interaction with these unrelated proteins therefore emerges from the optimization rather than from explicit counter-selection against them. Together, these results show that site-directed, dual-action peptides can selectively weaken a condensation hot spot, disrupt the native CHERP interaction condensate network, and destabilize the biomolecular condensate while maintaining low predicted cross-reactivity against unrelated test proteins.

## DISCUSSION

Dynamic protein surfaces, intrinsically disordered regions, and biomolecular condensates remain difficult targets for conventional small-molecule discovery because their function is often governed by conformational heterogeneity and distributed interaction networks rather than stable binding pockets. Here, we introduced PepSpace, an automated, physics-driven directed-evolution platform that couples multi-objective genetic optimization with residue-resolution coarse-grained molecular dynamics simulations. By evaluating peptide candidates across conformational ensembles, PepSpace directly incorporates molecular flexibility, multivalent interactions, solution conditions, self-association, and off-target binding into the peptide design objective. This physical selection layer extends beyond the usual objective space of sequence- and structure-generative models, which do not explicitly evaluate these state-dependent interactions during sequence generation, and enables the design of peptides that selectively and effectively engage dynamic protein targets whose recognition cannot be captured by a single static structure.

Across three demanding validation settings spanning structured, disordered, and condensed protein states, PepSpace showed consistent and transferable design performance. First, we benchmarked PepSpace against the generative masked language model *pepMLM* and the structural diffusion model *RFdiffusion* using the extracellular domains of AMHR2 and NCAM1. PepSpace-designed peptides achieved predicted interaction strengths up to one to two orders of magnitude greater than the machine-learning baselines, while maintaining high sequence diversity and discovering alternative binding modes not reached by the generative models. Unconstrained evolution further showed that affinity-focused optimization readily exploits promiscuous hydrophobic and aromatic surface patches, highlighting the importance of explicit negative selection for achieving target selectivity. Second, on the 2,500-residue mechanosensitive ion channel PIEZO1, PepSpace evolved 25-mer peptides that preferentially engage a defined 30-residue intracellular beam epitope while suppressing interactions across the remainder of the protein by more than an order of magnitude. Third, PepSpace enabled the *de novo* design of dual-action peptides targeting the condensation hot spot of the oncogenic splicing factor CHERP. By combining strong hotspot engagement with penalties against peptide self-association, the resulting sequences weaken native CHERP–CHERP interactions and destabilize the condensed phase.

A key architectural strength of PepSpace is its modular design. The simulations reported here used the coarse-grained Mpipi-Recharged force field^52^ implemented in LAMMPS^56^, which provides the sampling efficiency required for iterative, microsecond-scale population screening. Importantly, the directed-evolution framework itself is implemented independently of the underlying interaction potential and can be coupled to alternative simulation engines that provide compatible energetic and contact-based observables. These include coarse-grained models such as HPS^49,77^, CALVADOS^78,79^, and Martini3^80^, as well as atomistic explicit-solvent force fields such as CHARMM36m^81^ and AMBERff19SB^82^, implemented in engines including GROMACS^83^, OpenMM^84^, or NAMD^85^. This modularity supports a hierarchical design strategy in which broad sequence exploration is performed efficiently at coarse-grained resolution, followed by higher-resolution refinement of prioritized candidates where additional structural detail is required.

Taken together, PepSpace provides a versatile computational framework for designing selective peptide binders against dynamic protein interfaces and biomolecular condensates^29–32,86^. Future work will integrate active-learning surrogate models to accelerate sequence exploration^87^, introduce all-atom simulations to resolve binding kinetics, residence times, and off-rates, and evaluate selected leads in cellular systems. Coupling PepSpace-designed peptides to cell-penetrating or other delivery strategies may further enable intracellular and *in vivo* applications^88^.

## METHODS

### Residue-Resolution Coarse-Grained Molecular Dynamics Model

All coarse-grained molecular dynamics simulations were conducted using the Mpipi-Recharged force field^52^ executed in the LAMMPS simulation engine^56^. In this model, each amino acid residue is represented as a single interaction site located at the C*α* position with its characteristic mass, van der Waals radius *σ*_*i*_, and formal charge *q*_*i*_. Solvent dielectric screening and hydrophobic driving forces are modeled implicitly via effective residue pair potentials, enabling microsecond-scale conformational sampling. Consecutive residues along polypeptide backbones are connected via harmonic bonds (*r*_0_ = 3.81 Å, *k*_*b*_ = 8033.3 kJ/(mol *·* Å^2^)). Non-bonded interactions combine short-range contacts modeled by the Wang-Frenkel (WF) potential^89^ and electrostatic forces governed by an asymmetric screened Yukawa potential that captures side-chain reorientation and associative salt-bridge formation. Rigid globular domain boundaries for structured regions (AMHR2, NCAM1, PIEZO1, and CHERP) were assigned based on AlphaFold coordinates and experimental structures (SM Section SI, *Target Proteins*). Complete mathematical potential formulations, asymmetric Yukawa parameters, formal residue charges, and the full 20 *×* 20 Wang-Frenkel interaction parameter matrix are detailed in SM Section SII (Equations S1–S6 and Tables S2–S4).

### Simulation Protocols and Engine

All molecular dynamics simulations were performed using the LAMMPS molecular dynamics package^56^ compiled with custom pair styles implementing the Mpipi-Recharged potential. Langevin dynamics was employed to maintain constant temperature with a damping parameter of *γ* = 5000 fs (or 5.0 ps). An integration time step of Δ*t* = 10 fs was utilized across all production simulations following initial energy minimization and multi-stage timestep thermalization (0.5–7.0 fs). All simulations were conducted at an electrolyte ionic strength of 150 mM NaCl (*c*_salt_ = 150 mM, corresponding to a temperature-dependent Debye screening length of *κ*^−1^ *≈* 7.9 Å at 290 K).

### Evolutionary Algorithm Simulation Protocol

During directed evolution optimization cycles, candidate peptides and target proteins were instantiated in cubic simulation boxes with periodic boundary conditions and implicit aqueous solvent at *T* = 290 K and 150 mM NaCl. For AMHR2 and NCAM1 extracellular domain simulations, the box dimensions were set to 180*×*180*×*180 Å^3^, containing 1 target protein and 5 copies of the candidate peptide. For PIEZO1 monomer simulations, the box dimensions were set to 260 *×* 260 *×* 260 Å^3^, containing 1 full-length PIEZO1 monomer (2547 residues) and 5 peptide copies. Each generation evaluated a population of 15 candidate peptides over 50 evolutionary rounds. For each candidate peptide in every generation, two independent physical simulations were conducted: (1) a target-binding simulation to evaluate target affinity (*f*_bind_) and off-target cross-reactivity (*f*_cross_), and (2) a multi-copy peptide-only simulation in solvent to evaluate homotypic self-aggregation propensity (*f*_agg_), totaling 30 distinct physical simulations per generation. In each round, each simulation was integrated for 4 *×* 10^6^ MD steps (*τ* = 40 ns of aggregate conformational sampling per trajectory) to balance rapid evolutionary throughput with reliable sampling of initial binding pathways. Each simulation was executed using 16 MPI tasks (16 cores per simulation job). When submitted concurrently across cluster nodes in SLURM (utilizing 15 *×* 2 *×* 16 = 480 parallel MPI tasks per generation), each generation required *∼* 19 minutes of wall-clock time, and the full 50-round evolutionary campaign (comprising 1,500 independent trajectories: 750 binding and 750 self-aggregation simulations) completed in approximately 16 hours of aggregate wall-clock time. Users can freely select the number of MPI tasks per simulation and configure job distribution across cluster nodes to tailor wall-clock throughput to specific computational resources and queue allocations.

### Long-Time Production Simulations for Comparative Leads

To assess the equilibrium thermodynamic stability, contact persistence, and interaction networks of top-scoring leads and comparative designs outside the evolutionary optimization loop, extensive independent production runs were conducted for the prioritized candidates. For AMHR2 and NCAM1 extracellular domain systems, simulations were conducted in 180 *×* 180 *×* 180 Å^3^ cubic periodic boxes containing 1 target receptor and 5 peptide copies at *T* = 290 K and 150 mM NaCl for 6 *×* 10^8^ steps (6.0 *µ*s of aggregate sampling per candidate) across cognate, ML-designed, and cross-reactive pairs. Statistical uncertainty was quantified through trajectory block-averaging (SM Section SIII, Equations S7–S10). Detailed comparative statistics, sequence diversity dynamics, and domain-resolved binding decompositions are documented in SM Section SIV, SM Tables S5–S8, and SM Figures S1 and S2.

### Long-Time PIEZO1 Beam Targeting Simulations

To adequately sample the vast conformational space of the 2547-residue channel monomer and confirm sustained site-specific anchoring without off-target drift, long-time independent production simulations were conducted for the prioritized designs (*PIEZO1-01* and *PIEZO1-02*). Simulations were instantiated in cubic periodic boxes (260 *×* 260 *×* 260 Å^3^) containing 1 full-length PIEZO1 monomer and 5 candidate peptide copies in implicit aqueous electrolyte at *T* = 290 K and 150 mM NaCl without an explicit lipid bilayer for 2 *×* 10^9^ MD steps (20.0 *µ*s of aggregate sampling), focusing on the solvated intracellular beam lever loop while maintaining the rigid transmembrane and blade domains as an extensive steric and hydrophobic off-target counter-selection landscape. These production trajectories confirmed persistent spatial localization onto the designated beam epitope without non-specific adhesion across the surrounding channel body. Inter-replicate convergence, epitope specificity metrics, and whole-channel cross-reactivity profiling are detailed in SM Section SIV and SM Tables S9–S10.

### CHERP Condensate Direct Coexistence Simulations

For CHERP biomolecular condensates, homotypic liquid-liquid phase separation and its modulation by candidate peptides were characterized using direct coexistence (slab) simulations in elongated orthorhombic boxes (150 *×* 150 *×* 600 Å^3^) containing 25 full-length CHERP chains (916 residues each) and 150 mM NaCl^60,70^. For peptide-treated systems, 50 peptide molecules were introduced into the coexistence box. Direct coexistence simulations were equilibrated and produced over 2 *×* 10^8^ steps (2.0 *µ*s) per state point across temperatures ranging from 300–400 K. The equilibrium coexistence densities *ρ*_dense_(*T*) and *ρ*_dilute_(*T*) were extracted from the time-averaged 1D slab density profiles along the elongated *z*-axis. The critical solution temperature (*T*_*c*_) was determined by fitting the coexistence densities to the law of rectilinear diameters and 3D Ising critical scaling:

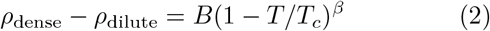

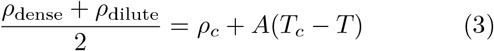

with the 3D Ising critical exponent *β* = 0.326^70^. Detailed residue-level contact and energetic profiling for CHERP homotypic interactions and peptide displacement were evaluated at *T* = 370 K near the critical phase boundary. Critical scaling regression covariance, fitted critical parameters (*T*_*c*_, *ρ*_*c*_), and homotypic contact attenuation statistics are detailed in SM Sections SIII and SIV, Equations S11–S14, and SM Tables S11–S13.

### Directed Evolution Pipeline, Genetic Operators, and Sequence Constraints

The genetic algorithm manages a population of candidate peptides across successive generations (*N* = 15 candidates per generation). Full amino acid sequences, physicochemical parameters (net formal charge, GRAVY, and polar fraction), and structural classifications for all generated and benchmark peptides are provided in SM Section SI and SM Table S1. To guarantee therapeutic developability and prevent the emergence of insoluble, hyper-sticky, or immunogenic sequences, candidate exploration is governed by multi-parametric sequence constraints enforced during initial population generation and every mutation/crossover event: **Net formal charge modulation**. The total net formal charge *q*_net_ = ∑_*i*_*q*_*i*_ (counting Arg/Lys as +1, Asp/Glu as −1, and His as +0.5) is restricted to the interval [−2.0, +5.0], preventing extreme electrostatic repulsion or non-specific polycationic cell toxicity while permitting physiological salt-bridge formation.

#### Hydropathy and solubility control

The Grand Average of Hydropathicity (GRAVY score) is maintained within [−1.5, +1.0], balancing hydrophobic contact capability with water solubility. Furthermore, the overall proportion of polar and charged residues (Arg, Lys, His, Asp, Glu, Ser, Thr, Asn, Gln) is constrained between 40% and 100% of the peptide sequence, ensuring robust aqueous solubility.

#### Cluster filtering capabilities

The platform additionally allows users to penalize or restrict clusters of contiguous residues (such as runs of identical amino acids or hydrophobic stretches), preventing kinetic aggregation and fibril nucleation when required by the design target.

#### Aromatic and oxidizable residue limits

Bulky aromatic and easily oxidized residues are explicitly regulated to preserve developability: each aromatic residue type (Trp, Phe, Tyr) is capped at *≤* 20% of the peptide length, and reactive residues such as Met are minimized, mitigating potential immunogenic epitopes, chemical degradation, and non-specific entropic adhesion. Offspring sequences adhering to these filters are generated using two primary genetic operators:

#### Contact-aware recombination

Parent pairs are selected from the top 50% of the population via tournament selection. Crossover breakpoints are weighted by per-residue interaction scores, ensuring that continuous contiguous peptide blocks that exhibit favorable interaction energies with the target are preferentially preserved.

#### Dynamic adaptive mutation

Point mutations are introduced with a baseline per-residue mutation probability *p*_mut_ = 0.15. Residues exhibiting strong, persistent target contacts (*E*_res_ *< −* 0.5 kcal/mol) have their mutation probability scaled down by a factor of 4 to conserve critical binding motifs, whereas non-contacting or repulsive residues have their mutation probability scaled up by 2 to encourage exploratory sequence variation, with mutated offspring validated against the developability constraints.

### Trajectory Analysis and Energetic Metrics

Intermolecular contact maps and interaction energies were computed using custom Python analysis pipelines leveraging MDAnalysis and NumPy. Two residues *i* and *j* were considered in physical contact if their C*α*–C*α* distance satisfied *r*_*ij*_ *<* 1.25*σ*_*ij*_. The pairwise interaction energy *E*_*ij*_ was calculated from the sum of non-bonded Lennard-Jones/Wang-Frenkel and screened Yukawa potentials averaged across all sampled trajectory frames. Statistical uncertainty quantification and standard errors of the mean (SEM) for all trajectory-averaged observables were computed via block-averaging (SM Section SIII, Equations S7–S10).

To ensure rigorous physical comparability across systems with differing chain lengths, copy numbers, and simulation geometries, observables are normalized as follows:

#### Peptide–Target Binding Energy per Peptide Residue (Figs. 2 and 3)

In multi-copy peptide binding assays (*N*_target_ = 1 chain, *N*_pep_ = 5 chains, each of length *L*_pep_), the total intermolecular interaction energy between target and peptide molecules *E*_inter_ is computed across all *N*_shots_ frames and *N*_target_ *· N*_pep_ molecular pairs:

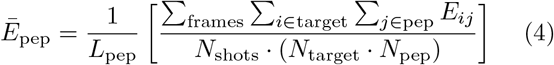

Normalizing by *L*_pep_ yields the mean binding interaction energy per amino acid (kcal/mol per residue), enabling unbiased interaction energy comparisons between peptides of different lengths (e.g., 12-mers versus 25-mers).

#### Whole-Protein Per-Residue Profiles (Figs. 3c,d, 4c,d, and 5d)

The local binding energy experienced by residue *i* of target protein *α* interacting with partner molecule *β* is computed as:

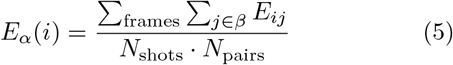

For the PIEZO1 monomer (Fig. 4c,d), values are integrated over 50-residue contiguous sliding windows (*w* = 50) to evaluate the macroscopic distribution of binding intensity along the 2547-residue polypeptide chain.

#### Multichain Condensate Interaction Energy per Pair (Fig. 5f,g)

In the 25-chain CHERP condensate simulations (*N*_*c*_ = 25 chains, *N*_*c*_(*N*_*c*_ −1)*/*2 = 300 intermolecular pairs), the homotypic interaction energy per CHERP–CHERP molecular pair is evaluated as:

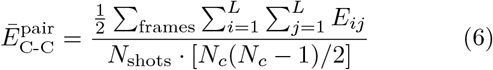

which computes the mean intermolecular contact interaction energy per CHERP–CHERP macromolecular pair across all 300 intermolecular pairs in the condensate ensemble. For the condensation hot spot (Fig. 5g), the summation is restricted strictly to residues *i, j ∈* [476, 576], quantifying the fraction of the total pairwise interaction energy localized solely to hot-spot self-interactions. For the per-residue interaction energy profile along the CHERP sequence (Fig. 5d), the local interaction energy of residue is evaluated as 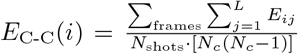 (in units of kcal/mol per residue per pair), which directly satisfies 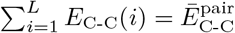.

## ACKNOWLEDGMENTS

J.O. acknowleges funding from CRIS Cancer Foundation for the research grant CRIS-CANCER-4332687. A. R. T. acknowledges funding from Ministerio de Ciencia e Innovacion under the Juan de la Cierva fellowship (JDC2024-053759-I). A. F. acknowledges funding from the Ramon y Cajal fellowship (RYC2021-030937-I) and Spanish National Grant (PID2022-136919NA-C33). A.O. acknowledges funding from CRIS Cancer Foundation (AOF.C01CRIS and AOF.M01CRIS). J.R.E. acknowledges funding from the Ramon y Cajal fellowship (RYC2021-030937-I), the Spanish National Agency for Research (PID2022-136919NA-C33 and PID2025-169417NB-C21), and the European Research Council (ERC) under the European Union’s Horizon Europe research and innovation program (grant agreement no. 101160499). J.R.E also acknowledges the CRIS Cancer Foundation for the research grant CRIS-CANCER-4332687. The authors acknowledge the computational resources provided by the Red Española de Supercomputación (RES) at the Barcelona Supercomputing Center (BSC), through projects FI-2025-3-0003, FI-2025-3-0065 and EHPC-REG-2025R02-173 on the MareNostrum5 supercomputer, and the computational resources at the CIEMAT Xula super-computer through project FI-2026-1-0027.

