## Supplementary Information for "PepSpace: An Automated, Physics-Driven Directed Evolution Platform for the De Novo Design of Specific Peptide Binders and Condensate Modulators"

Andrés R. Tejedor

*Department of Physical Chemistry,*
*Universidad Complutense de Madrid, 28040 Madrid, Spain*
*Instituto Pluridisciplinar, Universidad Complutense de Madrid, 28040 Madrid, Spain and*
*Yusuf Hamied Department of Chemistry, University of Cambridge,*
*Lensfield Road, Cambridge CB2 1EW, UK*

Maria Velasco-Estevez

*H12O-CNIO Hematological Malignancies Clinical Research Unit,*
*Spanish National Cancer Research Centre (CNIO), Madrid 28029, Spain*

Alberto Ocaña\*

*Experimental Therapeutics Unit, Hospital Clínico San Carlos (HCSC),*
*Instituto de Investigación Sanitaria San Carlos (IdISSC),*
*and CIBERONC, Madrid, Spain and*
*PhAsIca Biosciences S.L., Madrid, Spain*

Rosana Collepardo-Guevara†

*Yusuf Hamied Department of Chemistry, University of Cambridge,*
*Lensfield Road, Cambridge CB2 1EW, UK*
*Department of Genetics, University of Cambridge,*
*Downing Street, Cambridge CB2 3EH, UK and*

*PhAsIca Biosciences S.L., Madrid, Spain*

Jorge R. Espinosa<sup>‡</sup>

*Department of Physical Chemistry,*

*Universidad Complutense de Madrid, 28040 Madrid, Spain*

*Instituto Pluridisciplinar, Universidad Complutense de Madrid, 28040 Madrid, Spain*

*Yusuf Hamied Department of Chemistry, University of Cambridge,*

*Lensfield Road, Cambridge CB2 1EW, UK and*

*PhAsIca Biosciences S.L., Madrid, Spain*

(Dated: September 18, 2026)

\*

†

‡

### SI. SEQUENCES AND STRUCTURAL ARCHITECTURES OF MODELED PROTEINS AND PEPTIDES

Here, we provide the complete amino acid sequences, structural models, and rigid globular domain partitioning corresponding to all target proteins and candidate peptides investigated and plotted in the main manuscript and supplementary analyses.

#### A. Target Proteins

In our coarse-grained molecular dynamics simulations using the Mpipi-Recharged model, structured domains are treated as rigid bodies according to the AlphaFold predicted coordinates or experimental structures, while intrinsically disordered regions (IDRs) and flexible inter-domain linkers remain fully flexible with harmonic bonded connectivity ($r_0 = 3.8 \text{ \AA}$, $k_b = 8033.3 \text{ kJ}/(\text{mol} \cdot \text{\AA}^2)$). The rigid domain definitions are explicitly implemented in the LAMMPS simulation inputs via atom group commands (*group CD\** or *group rigid id ...*), where atom IDs corresponding to the first monomer chain define the globular structured segments. All coarse-grained simulations employ an implicit aqueous electrolyte representation (150 mM NaCl), wherein dielectric screening and effective hydrophobic driving forces are parameterized through the screened Yukawa and Wang-Frenkel pair potentials without explicit water molecules. For membrane-associated targets such as the 2547-residue PIEZO1 monomer, the system is simulated in this implicit aqueous environment without an explicit lipid bilayer, deliberately targeting the solvent-exposed, intracellular cytosolic lever-arm beam loop (residues 1373–1402) while retaining the extensive transmembrane blades and pore as a whole-channel steric and hydrophobic competitor for off-target counter-selection.

##### AMHR2 Extracellular Domain (128 residues):

PPNRRTCVFF EAPGVRGSTK TLGELLDTGT ELPRAIRCLY SRCCFGIWNL TQDRAQVEMQ GCRDSDEPGC
ESLHCDPSPR AHPSPGSTLF TCSCGTDFCN ANYSHLPPPG SPGTPGSQGP QAAPGESI

The structured extracellular domain of Anti-Müllerian Hormone Receptor Type 2 (AMHR2) was modeled based on the AlphaFold structural prediction. In the simulation input scripts, the rigid globular segments of the monomer (residues 1–128) are defined by:

**AMHR2 Rigid Residues:** 4–14, 19–26, 34–52, 54–65, 70–80, and 86–103.

The remaining terminal regions and loop segments (residues 1–3, 15–18, 27–33, 53, 66–69,

81–85, and 104–128) are treated as flexible chain regions.

**NCAM1 Extracellular Receptor (709 residues):**

```
MLQTKDLIWT LFFLGTAVSL QVDIVPSQGE ISVGESKFFL CQVAGDAKDK DISWFSPNGE KLTPNQQRIS
VWVNDSSST LTIYNANIDD AGIYKCVVTG EDGSESEATV NVKIFQKLMF KNAPTPQEFR EGEDAVIVCD
VVSSLPPTII WKHKGRDVIL KKDVRFIVLS NNYLQIRGIK KTDEGTYRCE GRILARGEIN FKDIQVIVNV
PPTIQARQNI VNATANLGQS VTLVCDAEGF PEPTMSWTKD GEQIEQEEDD EKYIFSDDSS QLTIKKVDKN
DEAEYICIAE NKAGEQDATI HLKVFAKPKI TYVENQTAME LEEQVTLTCE ASGDPIPSIT WRTSTRNISS
EEKASWTRPE KQETLDGHMV VRSHARVSSL TLKSIQYTDA GEYICTASNT IGQDSQSMYL EVQYAPKLQG
PVAVYTWEGN QVNITCEVFA YPSATISWFR DGQLLPSSNY SNIKIYNTPS ASYLEVTPDS ENDFGNYNCT
AVNRIGQESL EFILVQADTP SSPSIDQVEP YSSTAQVQFD EPEATGGVPI LKYKAEWRAV GEEVWHSKWY
DAKEASMEGI VTIVGLKPET TYAVRLAALN GKGLGEISAA SEFKTQPVQG EPSAPKLEGQ MGEDGNSIKV
NLIKQDDGGS PIRHYLVRYR ALSSEWKPEI RLPSGSDHVM LKSLDWNAEY EVYVVAENQQ GKSKAAHFVF
RTSAQPTAI
```

Neural Cell Adhesion Molecule 1 (NCAM1) is a multi-domain cell-surface receptor comprising five immunoglobulin-like (Ig1–Ig5) domains and two fibronectin type III (FnIII) repeats. In the multi-domain simulations, the rigid cores of the Ig and FnIII domains are partitioned into two main structural clusters: **NCAM1 Rigid Group CD1 (Ig1–Ig3 domains):** Residues 23–26, 36–41, 53–58, 79–84, 90–97, 107–114, 122–141, 149–153, 165–169, 173–193, 199–222, 230–254, 262–266, 268–275, 280–318, and 325–333.

**NCAM1 Rigid Group CD2 (Ig4–FnIII domains):** Residues 377–382, 391–399, 402–450, 462–468, 470–480, 483–520, 525–550, 554–563, 569–573, 580–604, 615–621, 627–632, 644–651, 655–662, 668–672, 678–687, and 692–702.

All inter-domain hinges, loop insertions, and terminal tails remain completely flexible.

**PIEZO1 Monomer (2521 residues in modeled coordinate frame):**

```
MEPHVLGAVL YWLLLPALL AACLLRFSGS SLVYLLFLLL LPWFPGPTRC GLWLLLRASL VLLGPLAWLL
LLALPSWPRP PRPQGTGPGSV AASLLLLLLL LCRCAPCLQ LWLALALLTG AAELFWLLF RVWDQPFLLR
LLHWVAAGAL LLLGAPCLAL LPRLRWVLE LCGALLPRLS LALLLWPLVL LALRALLPG PRPLALLRFL
ALLLALLSL LPWPRTGPLV LLLLWLLLLC HLLCGLRALQ LPALWPGFLF LCRACPVLLE AWLALCALLL
LRALVLLALQ CGLRPALRAL LPLLLLLLWG ALLALPVFLF LLLRPLALLL PLLLPWSQL LLLLPWPLAL
LLGLPCLALQ LPALPWLLLL LLPLALLCAL LLRWPLGLL LLLLPLVLLL LPSLGLLLL LPVLLLPLG
LLLLPLVLL LPALLLLLLP PVLLLLPPLP WLLLLPPLAL LLLLPLPWL LLLLPLPW LLLLPLPW
LLLLPLPWL LLLLPLPWL LLLLPLPWL LLLLPLPWL LLLLPLPWL LLLLPLPWL LLLLPLPWL
LLLLPLALL LLLPLALL LLLPLALL LPLALLLL LPLALLLL LPLALLLL LPLALLLL LPLALLLL
ALPLPLPLA LLLLPLPLA LLLLPLPLA LLLPLALL LLLPLALL LPLALLLL LPLALLLL LPLALLLL
PLALLLLLP LALLLLLP LALLLLLP LLLPLPLA LLLPLPLA LLLPLPLA LLLPLPLA LLLPLPLA
LPPLALLLL PPLALLLL PPLALLLL LALLLLLP LALLLLLP LLLPLPLA LLLPLPLA LLLPLPLA
LLPLALLL LLLPLALL LLLPLALL LLLPLALL LLLPLALL LLLPLALL LLLPLALL LLLPLALL
LALLLLLP LLLPLALL LLLPLALL LLLPLALL LLLPLALL LLLPLALL LLLPLALL LLLPLALL
PPLALLLL LLLPLALL LLLPLALL LLLPLALL LLLPLALL LLLPLALL LLLPLALL LLLPLALL
LLPLALLL LPLALLLL LPLALLLL LPLALLLL LPLALLLL LPLALLLL LPLALLLL LPLALLLL
LLPLALLL LLLPLALL LLLPLALL LLLPLALL LLLPLALL LLLPLALL LLLPLALL LLLPLALL
ALLLLPLA LLLPLPLA LLLPLPLA LLLPLALL LLLPLALL LLLPLALL LLLPLALL LLLPLALL
```

Human mechanosensitive channel PIEZO1 forms a massive curved trimer of 38 transmembrane-helix blades and a central intracellular helical beam. The single monomer modeled in this work comprises the propeller blade helices and beam lever arm (AlphaFold ID: AF-Q92508-F1). In the simulation inputs, the rigid structural units of the single monomer are defined by:

**PIEZO1 Monomer Rigid Residues:** 11–20, 22–29, 31–41, 57–75, 123–126, 128–133, 201–238, 243–268, 270–276, 278–291, 309–327, 425–454, 466–488, 490–493, 506–511, 513–532, 570–620, 622–650, 652–676, 678–712, 789–890, 892–922, 924–956, 963–1072, 1077–1135, 1143–1264, 1285–1366, 1409–1421, 1463–1468, 1510–1556, 1564–1569, 1643–1764, 1766–1806, 1936–1952, 1956–1982, 2002–2193, 2203–2278, 2300–2313, 2316–2326, 2336–2342, 2352–2392, 2402–2412, 2414–2427, and 2441–2519.

Crucially, the designated target epitope on the intracellular beam (residues 1373–1402) resides completely within the flexible region between residue 1367 and 1408, permitting realistic conformational adaptation and induced-fit binding during peptide engagement.

**CHERP (916 residues):**

MEMPLPPDDQ ELRNVIDKLA QFVARNGPEF EKMTMEKQKD NPKFSFLFGG EFYSYYKCKL ALEQQQLICK
QQTPELEPAA TMPPLPQPPL APAAPIPPAQ GAPSMDELIQ QSQWNLQQQE QHLLALRQEQ VTA AVAHAVE
QQMQKLLEET QLDMNEFDNL LQPIIDTCTK DAISAGKNWM FSNAKSPPHC ELMAGHLRNR ITADGAHFEL
RLHLIYLIND VLHHCQRKQA RELLAALQKV VVPIYCTSFL AVEEDKQQKI ARLLQLWEKN GYFDDSIIQQ
LQSPALGLGQ YQATLINEYS SVVQPVQLAF QQQIQTLKTQ HEEFVTS LAQ QQQQQQQQQQ QLQMPQMEAE
VKATPPPPAP PPAPAPAPAI PPTTQPDDSK PPIQMPGSSE YEAPGGVQDP AAAGPRGPGP HDQIPPNKPP
WFDQPHPVAP WGQQQPPEQP PYPHHQGGPP HCPPWNNSHE GMWGEQRGDP GWNGQRDAPW NNQPDAAWNS
QFEGPWNSQH EQPPWGGGQR EPPFRMQRPP HFRGPFPPHQ QHPQFNQPPH PHNFNRFPPR FMQDDFPPRH
PFERPPYPHR FDYPQGD FPA EMGPPHHHPG HRMPHPGINE HPPWAGPQHP DFGPPPHGFN GQPPHMRRQG
PPHINHDDPS LVPNVPYFDL PAGLMAPLVK LEDHEYKPLD PKDIRLPPPM PPSERLLA AV EAFYSPPSHD
RPRNSEGWEQ NGLYEFFRAK MRARRRKGQE KRNSGPSRSR SRSKSRGRSS SRSNSRSSKS SGSYSRSRSR
SCSRSYSRSR SRSRSRSRSS RSRSRSQSRS RSKSYSPGRR RRSRSRSPTP PSSAGLGSNS APPIPDSRLG
EENKGHQMLV KMGWSGSGGL GAKEQGIQDP IKGGDVRDKW DQYKGVGVAL DDPYENYRRN KSYSFIARMK
ARDECK

Calcium Homeostasis Endoplasmic Reticulum Protein (CHERP) is an oncogenic RNA-processing and splicing factor consisting of an N-terminal structured helical fold and an extensive, multivalent low-complexity domain driving liquid-liquid phase separation (LLPS). In our multi-chain condensate simulations (25 chains), the structured core of CHERP (monomer range 1–916) is defined by two rigid groups: **CHERP Rigid Group CD1:** Residues 14–20.

**CHERP Rigid Group CD2:** Residues 124–202, 205–250, 256–270, and 272–326.

The remaining 65% of the CHERP sequence (residues 1–13, 21–123, 203–204, 251–255, 271, and 327–916) is entirely disordered and flexible. Notably, the critical intermolecular condensation hot spot (residues 476–576) discovered by PepSpace is entirely located within the flexible low-complexity IDR domain.

#### B. Candidate Peptides and Sequence Developability Constraints

Unlike the folded domains of multi-domain proteins, **all candidate and benchmark peptides utilized in this work are modeled as fully flexible linear polypeptides** ($r_0 = 3.81 \text{ \AA}$, $k_b = 8033.3 \text{ kJ}/(\text{mol}\cdot\text{\AA}^2)$). Peptides do not contain rigid bodies or internal constraints unless explicitly disulfide-cyclized.

Furthermore, during directed evolution in PepSpace, candidate peptide exploration is governed by a multi-parametric developability filter to ensure high solubility, chemical stability, and low immunogenic potential: **Formal Net Charge**. Allowed in the range $[-2.0, +5.0]$ to permit favorable electrostatic complementarity while preventing extreme cationic or anionic toxicity.

**Hydropathy (GRAVY)**. Maintained between $-1.5$ and $+1.0$, balancing hydrophobic binding capability with aqueous solubility.

**Polar/Charged Residue Content**. Enforced between 40% and 100% of the total sequence length.

**Cluster Filtering Capabilities**. The platform allows users to penalize or restrict clusters of contiguous residues (such as runs of identical amino acids or hydrophobic stretches) to prevent kinetic aggregation and fibril formation when desired.

**Aromatic and Oxidizable Residue Limits**. Each aromatic residue type (Trp, Phe, Tyr) is capped at $\leq 20\%$ of the peptide length, and reactive residues such as Met are minimized to prevent non-specific entropic adhesion, chemical instability, and immunogenic epitopes. Table S1 lists all candidate peptides shown in the plots and discussed in the primary text.

TABLE S1. **Sequences, design methodology, and target designations for all candidate peptides evaluated in this work (Table S1).** Listed are the peptide designation, design methodology or category, target protein, and full single-letter amino acid sequence. Primary experimental benchmark leads are highlighted in parentheses.

| Peptide Designation | Method / Class | Target Protein | Amino Acid Sequence |
| --- | --- | --- | --- |
| *Machine-Learning and Experimental Benchmarks (AMHR2 and NCAM1)* |  |  |  |
| AMHR2-pepMLM-01 | pepMLM | AMHR2 | LTRYTSLAAQGC |
| AMHR2-pepMLM-02 | pepMLM | AMHR2 | DTTLYSGFAQYG |
| AMHR2-pepMLM-03 (*pepMLM-1*) | pepMLM | AMHR2 | DESLRSFLAHYC |
| AMHR2-pepMLM-04 | pepMLM | AMHR2 | LETYRSGLAQYC |
| AMHR2-RFD-01 (*RFD-1*) | RFdiffusion | AMHR2 | APAVTGSILADL |
| AMHR2-RFD-02 | RFdiffusion | AMHR2 | SAAVTGSILADL |
| AMHR2-RFD-03 | RFdiffusion | AMHR2 | RSIRLSYPVELP |
| AMHR2-RFD-04 | RFdiffusion | AMHR2 | RSVRLVHPVELP |
| NCAM1-pepMLM-01 | pepMLM | NCAM1 | GKLPLPSLPCK |
| NCAM1-pepMLM-02 (*pepMLM-2*) | pepMLM | NCAM1 | GLGPSPVLPRC |
| NCAM1-pepMLM-03 | pepMLM | NCAM1 | GLGPLPVLPCK |
| NCAM1-pepMLM-04 | pepMLM | NCAM1 | HSLGQPLSPICSE |
| NCAM1-RFD-01 | RFdiffusion | NCAM1 | SLPIENIYIEA |
| NCAM1-RFD-02 (*RFD-2*) | RFdiffusion | NCAM1 | MKPIEVVYEKA |
| NCAM1-RFD-03 | RFdiffusion | NCAM1 | ELPEQVIYIEA |
| NCAM1-RFD-04 | RFdiffusion | NCAM1 | EKPIEVIYEKA |
| *Top-5 Directed Evolution Leads (AMHR2 and NCAM1)* |  |  |  |
| top-01 AMHR2 | PepSpace | AMHR2 | CNKYSPSYWKFW |
| top-02 AMHR2 | PepSpace | AMHR2 | PQKTSLIYWKFW |
| top-03 AMHR2 | PepSpace | AMHR2 | YQKTSLICWKYW |
| top-04 AMHR2 | PepSpace | AMHR2 | EQKTSLSYWKFW |
| top-05 AMHR2 | PepSpace | AMHR2 | INGMHAQKWKYW |
| top-01 NCAM1 | PepSpace | NCAM1 | NIKFWRHWRIH |
| top-02 NCAM1 | PepSpace | NCAM1 | CKKFHV FYRWH |
| top-03 NCAM1 | PepSpace | NCAM1 | HKRWQYFKRVF |
| top-04 NCAM1 | PepSpace | NCAM1 | REKFHVHWRCW |
| top-05 NCAM1 | PepSpace | NCAM1 | FQRWQYFHRVE |
| *PIEZO1 Helical Beam Binders* |  |  |  |
| PIEZO1-01 | PepSpace | PIEZO1 (Beam) | HLQHSHFLDVERIRSPNRYNLRFSW |
| PIEZO1-02 | PepSpace | PIEZO1 (Beam) | DARQSPAAPPFKYYFSSYARREFYT |
| PIEZO1-03 | PepSpace | PIEZO1 (Beam) | CHPSKSMDSIWEAKTFWEP SHDHAL |
| *CHERP Condensate Modulators* |  |  |  |
| CHERP-01 | PepSpace | CHERP (Hotspot) | GMRMDLQHCLENLRFPRKTWRHSWKHF |
| CHERP-02 | PepSpace | CHERP (Hotspot) | YCHLTHSWDHRTWETCHVLFIHYSI |

#### SII. FORMULATION AND PARAMETRIZATION OF THE MPIPI-RECHARGED MODEL

The Mpipi-Recharged model [1] is an advanced residue-resolution coarse-grained force field designed to accurately capture the delicate balance of associative and repulsive forces governing biomolecular condensation, conformational ensembles of intrinsically disordered proteins (IDPs), and multivalent protein–protein and protein–peptide interactions.

##### A. Physical Origin and Charge Asymmetry

Standard coarse-grained protein models treat electrostatic interactions using symmetric screened Coulomb or Yukawa potentials, where like-charge repulsions (e.g., Arg–Arg, Glu–Glu) and opposite-charge attractions (e.g., Arg–Glu, Lys–Asp) share identical coupling magnitudes. However, explicit-solvent atomistic potential of mean force (PMF) calculations reveal a fundamental physical asymmetry: **Side-chain reorientation**. When like charges approach, conformational entropy penalty and side-chain rotamer flexibility allow charged moieties to reorient away from each other, effectively softening the repulsive barrier compared to rigid point-charge Coulomb’s law.

**Associative ion pairing and hydration.** When opposite charges approach, water structure and directional salt-bridge formation produce strong, associative free-energy minima that significantly exceed the magnitude of like-charge repulsions at short distances. Mpipi-Recharged addresses this asymmetry by decoupling the short-range contact potential from the electrostatic interactions and introducing an asymmetric pair-specific Yukawa interaction parameter $A_{ij}$ calibrated from PMF calculations and liquid-liquid phase separation benchmarks.

##### B. Potential Energy Functions

In the Mpipi-Recharged force field, bonded connectivity along the chain backbone is represented by harmonic springs:

$$V_{\text{bond}}(r) = \frac{k_b}{2}(r - r_0)^2 \tag{S1}$$

where $k_b = 8033.3 \text{ kJ}/(\text{mol} \cdot \text{\AA}^2)$ ($9.600 \text{ kcal}/(\text{mol} \cdot \text{\AA}^2)$) and $r_0 = 3.81 \text{ \AA}$.

Non-bonded interactions consist of two independent additive contributions:

$$V_{\text{non-bonded}}(r) = V_{\text{WF}}(r) + V_{\text{Yukawa}}(r) \quad (\text{S2})$$

##### 1. Wang-Frenkel Short-Range Potential

Short-range non-electrostatic interactions (van der Waals, π-π stacking, cation-π, and hydrophobic interactions) are governed by the Wang-Frenkel (WF) potential [2]:

$$V_{\text{WF}}(r) = \epsilon_{ij} \alpha_{ij} \left[ \left( \frac{\sigma_{ij}}{r} \right)^{2\mu} - 1 \right] \left[ \left( \frac{r_{\text{cut},ij}}{r} \right)^{2\mu} - 1 \right]^{2\nu} \quad (\text{S3})$$

for $r \leq r_{\text{cut},ij}$, and $V_{\text{WF}}(r) = 0$ for $r > r_{\text{cut},ij}$, where the cutoff radius is defined as $r_{\text{cut},ij} = 3.0\sigma_{ij}$. The parameter $\alpha_{ij}$ is a normalization constant that guarantees that the minimum of the potential occurs exactly at depth $-\epsilon_{ij}$:

$$\alpha_{ij} = 2\nu \left( \frac{\sigma_{ij}}{r_{\text{cut},ij}} \right)^{2\mu} \left( \frac{1 + 2\nu}{2\nu [1 - (\sigma_{ij}/r_{\text{cut},ij})^{2\mu}] } \right)^{2\nu+1} \quad (\text{S4})$$

For interactions involving rigid globular regions, non-bonded interaction strengths are scaled to account for reduced surface accessibility:

$$\epsilon_{ij}^{\text{eff}} = \begin{cases} \epsilon_{ij}, & \text{IDP - IDP} \\ \sqrt{0.7} \epsilon_{ij}, & \text{IDP - Globular} \\ 0.7 \epsilon_{ij}, & \text{Globular - Globular} \end{cases} \quad (\text{S5})$$

##### 2. Asymmetric Yukawa Electrostatic Potential

Electrostatic interactions between charged amino acids ($i, j \in \{\text{R, K, H, D, E}\}$) are modeled via a screened Yukawa potential:

$$V_{\text{Yukawa}}(r) = A_{ij} \frac{\exp(-\kappa r)}{r} \quad (\text{S6})$$

where $\kappa$ is the inverse Debye screening length in Angstrom<sup>-1</sup>, calculated dynamically from the temperature $T$ and ionic strength $c_s$ (in mol/L or mM):

$$\kappa = \sqrt{\frac{8\pi l_B N_A c_s}{1000}} = \sqrt{8\pi l_B I} \quad (\text{S7})$$

Here, $l_B$ is the Bjerrum length:

$$l_B = \frac{e^2}{4\pi\epsilon_0\epsilon_r k_B T} \quad (\text{S8})$$

where the temperature-dependent relative dielectric permittivity of water $\epsilon_r(T)$ is evaluated using the Malmberg-Maryott polynomial:

$$\epsilon_r(T) = \frac{5321}{T} + 233.76 - 0.9297T + 0.001417T^2 - 0.0000008292T^3 \quad (\text{S9})$$

At $T = 290$ K and $c_s = 150$ mM NaCl, $\epsilon_r \approx 78.5$, yielding $l_B \approx 7.0$ Å and a Debye screening length of $\kappa^{-1} \approx 7.9$ Å.

The classical symmetric Coulombic relation is given by $E_{\text{Coulomb}} = \frac{k}{\epsilon_r} \frac{q_i q_j}{r}$, with $k = 1/4\pi\epsilon_0 = 331.61$ kcal·Å/(mol·e<sup>2</sup>). For a uniform medium with $\epsilon_r = 80$ and point charges $q_i = \pm 1e$, the coupling magnitude would be $|A| = 4.145$ kcal·Å/mol. In Mpipi-Recharged, the interaction parameter $A_{ij}$ is pair-specific, as tabulated in Table S2, and its equivalent effective charge product $q_i q_j$ is compiled in Table S3.

**TABLE S2. Parameter $A_{ij}$ (in kcal mol<sup>-1</sup> Å) for the asymmetric Yukawa electrostatic potential in the Mpipi-Recharged model (Table S2).**

|  | **R** | **K** | **H** | **D** | **E** |
| --- | --- | --- | --- | --- | --- |
| **R** | +4.00 | +4.00 | +1.47 | -4.91 | -4.93 |
| **K** | +4.00 | +4.00 | +1.48 | -4.33 | -4.34 |
| **H** | +1.47 | +1.48 | +1.04 | -2.46 | -2.45 |
| **D** | -4.91 | -4.33 | -2.46 | +4.00 | +4.00 |
| **E** | -4.93 | -4.34 | -2.45 | +4.00 | +4.00 |

Notice the significant physical asymmetry: attractive electrostatic pairings (e.g., Arg–Glu: −4.93, Arg–Asp: −4.91, Lys–Glu: −4.34) possess higher coupling magnitudes than like-charge repulsions (+4.00), accurately reproducing the excess associative free energy observed in explicit-solvent atomistic simulations while preserving the complete solubility of isolated poly-ampholyte sequences.

TABLE S3. **Effective charge product $q_i q_j$ equivalent to parameter $A_{ij}$ for the Yukawa potential in the Mpipi-Recharged model (Table S3).**

| | $q_R$ | $q_K$ | $q_H$ | $q_D$ | $q_E$ |
| --- | --- | --- | --- | --- | --- |
| $q_R$ | +0.963 | +0.963 | +0.354 | -1.182 | -1.187 |
| $q_K$ | +0.963 | +0.963 | +0.354 | -1.043 | -1.042 |
| $q_H$ | +0.354 | +0.354 | +0.250 | -0.593 | -0.592 |
| $q_D$ | -1.182 | -1.043 | -0.593 | +0.963 | +0.963 |
| $q_E$ | -1.187 | -1.042 | -0.592 | +0.963 | +0.963 |

##### C. Wang-Frenkel Interaction Matrix

The full set of non-bonded Wang-Frenkel parameters ($\epsilon_{ij}$, $\sigma_{ij}$, $\nu$, and $\mu$) for all canonical amino acids (including phosphorylated or non-standard residues where applicable) is compiled in Table S4.

TABLE S4: **Non-bonded interaction parameters for the Wang-Frenkel potential in the Mpipi-Recharged model (Table S4).**

| Residue $i^{\text{th}}$ | Residue $j^{\text{th}}$ | $\epsilon$ (kcal mol$^{-1}$) | $\sigma$ (Å) | $\nu$ | $\mu$ |
| --- | --- | --- | --- | --- | --- |
| **A** | **A** | 0.0912 | 5.2701 | 1 | 4 |
| **A** | **D** | 0.1363 | 5.5468 | 1 | 3 |
| **A** | **E** | 0.1437 | 5.7239 | 1 | 3 |
| **A** | **Y** | 0.3808 | 6.0019 | 1 | 3 |
| **A** | **V** | 0.0529 | 5.7680 | 1 | 4 |
| **A** | **L** | 0.0577 | 5.9021 | 1 | 4 |
| **A** | **Q** | 0.2226 | 5.7740 | 1 | 3 |
| **A** | **W** | 0.5005 | 6.1683 | 1 | 3 |
| **A** | **F** | 0.3582 | 5.9498 | 1 | 3 |
| **A** | **S** | 0.1017 | 5.3414 | 1 | 4 |
| **A** | **H** | 0.3680 | 5.8039 | 1 | 3 |
| **A** | **N** | 0.2169 | 5.5967 | 1 | 3 |
| **A** | **P** | 0.1166 | 5.5381 | 1 | 3 |
| **A** | **C** | 0.1084 | 5.4972 | 1 | 4 |
| **A** | **I** | 0.0484 | 6.0959 | 1 | 4 |

Table S4 continued from previous page

| Residue $i^{\text{th}}$ | Residue $j^{\text{th}}$ | $\epsilon$ (kcal mol$^{-1}$) | $\sigma$ (Å) | $\nu$ | $\mu$ |
| --- | --- | --- | --- | --- | --- |
| C | C | 0.1257 | 5.7244 | 1 | 3 |
| C | I | 0.0657 | 6.3230 | 1 | 4 |
| D | D | 0.1379 | 5.8235 | 1 | 3 |
| D | E | 0.1434 | 6.0006 | 1 | 3 |
| D | Y | 0.4132 | 6.2786 | 1 | 3 |
| D | V | 0.0994 | 6.0448 | 1 | 4 |
| D | L | 0.1040 | 6.1788 | 1 | 4 |
| D | Q | 0.2632 | 6.0507 | 1 | 3 |
| D | W | 0.5289 | 6.4450 | 1 | 3 |
| D | F | 0.3914 | 6.2265 | 1 | 3 |
| D | S | 0.1465 | 5.6181 | 1 | 3 |
| D | H | 0.0080 | 6.0807 | 1 | 7 |
| D | N | 0.2576 | 5.8734 | 1 | 3 |
| D | P | 0.1608 | 5.8148 | 1 | 3 |
| D | C | 0.1530 | 5.7739 | 1 | 3 |
| D | I | 0.0951 | 6.3726 | 1 | 4 |
| E | E | 0.1489 | 6.1777 | 1 | 3 |
| E | Y | 0.4202 | 6.4557 | 1 | 3 |
| E | V | 0.1068 | 6.2218 | 1 | 4 |
| E | L | 0.1114 | 6.3559 | 1 | 4 |
| E | Q | 0.2706 | 6.2278 | 1 | 3 |
| E | W | 0.5360 | 6.6221 | 1 | 3 |
| E | F | 0.3984 | 6.4036 | 1 | 3 |
| E | S | 0.1539 | 5.7952 | 1 | 3 |
| E | H | 0.0080 | 6.2577 | 1 | 7 |
| E | N | 0.2650 | 6.0505 | 1 | 3 |
| E | P | 0.1682 | 5.9919 | 1 | 3 |
| E | C | 0.1604 | 5.9510 | 1 | 3 |

Table S4 continued from previous page

| Residue $i^{\text{th}}$ | Residue $j^{\text{th}}$ | $\epsilon$ (kcal mol$^{-1}$) | $\sigma$ (Å) | $\nu$ | $\mu$ |
| --- | --- | --- | --- | --- | --- |
| **E** | **I** | 0.1025 | 6.5497 | 1 | 4 |
| **F** | **F** | 0.6004 | 6.6296 | 1 | 2 |
| **F** | **S** | 0.3681 | 6.0211 | 1 | 3 |
| **F** | **H** | 0.6198 | 6.4837 | 1 | 2 |
| **F** | **N** | 0.4770 | 6.2765 | 1 | 3 |
| **F** | **P** | 0.3822 | 6.2179 | 1 | 3 |
| **F** | **C** | 0.3745 | 6.1770 | 1 | 3 |
| **F** | **I** | 0.3178 | 6.7756 | 1 | 3 |
| **G** | **A** | 0.1321 | 4.9826 | 1 | 3 |
| **G** | **C** | 0.1493 | 5.2097 | 1 | 3 |
| **G** | **D** | 0.1758 | 5.2593 | 1 | 3 |
| **G** | **E** | 0.1832 | 5.4364 | 1 | 3 |
| **G** | **F** | 0.3968 | 5.6623 | 1 | 3 |
| **G** | **G** | 0.1730 | 4.6951 | 1 | 3 |
| **G** | **H** | 0.4089 | 5.5165 | 1 | 3 |
| **G** | **I** | 0.0893 | 5.8084 | 1 | 4 |
| **G** | **K** | 0.1231 | 5.6832 | 1 | 4 |
| **G** | **L** | 0.0986 | 5.6146 | 1 | 4 |
| **G** | **N** | 0.2578 | 5.3092 | 1 | 3 |
| **G** | **P** | 0.1575 | 5.2506 | 1 | 3 |
| **G** | **Q** | 0.2635 | 5.4865 | 1 | 3 |
| **G** | **R** | 0.3353 | 5.7671 | 1 | 3 |
| **G** | **S** | 0.1426 | 5.0539 | 1 | 3 |
| **G** | **T** | 0.1158 | 5.2921 | 1 | 3 |
| **G** | **V** | 0.0939 | 5.4806 | 1 | 4 |
| **G** | **W** | 0.5393 | 5.8808 | 1 | 3 |
| **G** | **Y** | 0.4195 | 5.7144 | 1 | 3 |
| **H** | **H** | 0.0524 | 6.3378 | 1 | 4 |

Table S4 continued from previous page

| Residue $i^{\text{th}}$ | Residue $j^{\text{th}}$ | $\epsilon$ (kcal mol$^{-1}$) | $\sigma$ (Å) | $\nu$ | $\mu$ |
| --- | --- | --- | --- | --- | --- |
| H | N | 0.4937 | 6.1306 | 1 | 3 |
| H | P | 0.3934 | 6.0720 | 1 | 3 |
| H | C | 0.3852 | 6.0311 | 1 | 3 |
| H | I | 0.3252 | 6.6297 | 1 | 3 |
| I | I | 0.0057 | 6.9217 | 1 | 12 |
| K | K | 0.0455 | 6.6713 | 1 | 5 |
| K | T | 0.0674 | 6.2802 | 1 | 4 |
| K | R | 0.2167 | 6.7552 | 1 | 3 |
| K | A | 0.0847 | 5.9707 | 1 | 4 |
| K | D | 0.0009 | 6.2474 | 1 | 9 |
| K | E | 0.0009 | 6.4245 | 1 | 9 |
| K | Y | 0.1842 | 6.7025 | 1 | 3 |
| K | V | 0.0442 | 6.4687 | 1 | 5 |
| K | L | 0.0492 | 6.6027 | 1 | 5 |
| K | Q | 0.2241 | 6.4746 | 1 | 3 |
| K | W | 0.1984 | 6.8690 | 1 | 3 |
| K | F | 0.2061 | 6.6504 | 1 | 3 |
| K | S | 0.0959 | 6.0420 | 1 | 4 |
| K | H | 0.1663 | 6.5046 | 1 | 3 |
| K | N | 0.2180 | 6.2973 | 1 | 3 |
| K | P | 0.1117 | 6.2388 | 1 | 4 |
| K | C | 0.1030 | 6.1979 | 1 | 4 |
| K | I | 0.0394 | 6.7965 | 1 | 5 |
| L | L | 0.0242 | 6.5341 | 1 | 6 |
| L | Q | 0.1891 | 6.4060 | 1 | 3 |
| L | W | 0.4687 | 6.8003 | 1 | 3 |
| L | F | 0.3265 | 6.5818 | 1 | 3 |
| L | S | 0.0682 | 5.9734 | 1 | 4 |

Table S4 continued from previous page

| Residue $i^{\text{th}}$ | Residue $j^{\text{th}}$ | $\epsilon$ (kcal mol$^{-1}$) | $\sigma$ (Å) | $\nu$ | $\mu$ |
| --- | --- | --- | --- | --- | --- |
| L | H | 0.3345 | 6.4359 | 1 | 3 |
| L | N | 0.1833 | 6.2287 | 1 | 3 |
| L | P | 0.0831 | 6.1701 | 1 | 4 |
| L | C | 0.0749 | 6.1292 | 1 | 4 |
| L | I | 0.0149 | 6.7279 | 1 | 6 |
| M | A | 0.0825 | 5.8690 | 1 | 4 |
| M | C | 0.0998 | 6.0962 | 1 | 4 |
| M | D | 0.1280 | 6.1457 | 1 | 3 |
| M | E | 0.1354 | 6.3228 | 1 | 3 |
| M | F | 0.3507 | 6.5488 | 1 | 3 |
| M | G | 0.1234 | 5.5815 | 1 | 3 |
| M | H | 0.3593 | 6.4029 | 1 | 3 |
| M | I | 0.0398 | 6.6948 | 1 | 5 |
| M | K | 0.0706 | 6.5696 | 1 | 4 |
| M | L | 0.0490 | 6.5010 | 1 | 4 |
| M | M | 0.0739 | 6.4680 | 1 | 4 |
| M | N | 0.2082 | 6.1957 | 1 | 3 |
| M | P | 0.1079 | 6.1371 | 1 | 4 |
| M | Q | 0.2140 | 6.3729 | 1 | 3 |
| M | R | 0.2876 | 6.6535 | 1 | 3 |
| M | S | 0.0931 | 5.9403 | 1 | 4 |
| M | T | 0.0662 | 6.1785 | 1 | 4 |
| M | V | 0.0443 | 6.3670 | 1 | 5 |
| M | W | 0.4923 | 6.7673 | 1 | 3 |
| M | Y | 0.3727 | 6.6008 | 1 | 3 |
| N | N | 0.3425 | 5.9234 | 1 | 3 |
| N | P | 0.2423 | 5.8648 | 1 | 3 |
| N | C | 0.2341 | 5.8239 | 1 | 3 |

Table S4 continued from previous page

| Residue $i^{\text{th}}$ | Residue $j^{\text{th}}$ | $\epsilon$ (kcal mol$^{-1}$) | $\sigma$ (Å) | $\nu$ | $\mu$ |
| --- | --- | --- | --- | --- | --- |
| N | I | 0.1741 | 6.4225 | 1 | 3 |
| P | P | 0.1420 | 5.8062 | 1 | 3 |
| P | C | 0.1338 | 5.7653 | 1 | 3 |
| P | I | 0.0738 | 6.3639 | 1 | 4 |
| Q | Q | 0.3540 | 6.2779 | 1 | 3 |
| Q | W | 0.6252 | 6.6722 | 1 | 2 |
| Q | F | 0.4824 | 6.4537 | 1 | 3 |
| Q | S | 0.2332 | 5.8453 | 1 | 3 |
| Q | H | 0.4994 | 6.3078 | 1 | 3 |
| Q | N | 0.3483 | 6.1006 | 1 | 3 |
| Q | P | 0.2480 | 6.0420 | 1 | 3 |
| Q | C | 0.2399 | 6.0011 | 1 | 3 |
| Q | I | 0.1799 | 6.5998 | 1 | 3 |
| R | R | 0.1507 | 6.8391 | 1 | 3 |
| R | A | 0.2959 | 6.0546 | 1 | 3 |
| R | D | 0.0067 | 6.3313 | 1 | 7 |
| R | E | 0.0069 | 6.5084 | 1 | 7 |
| R | Y | 0.8042 | 6.7863 | 1 | 2 |
| R | V | 0.2591 | 6.5525 | 1 | 3 |
| R | L | 0.2636 | 6.6866 | 1 | 3 |
| R | Q | 0.4225 | 6.5585 | 1 | 3 |
| R | W | 0.9341 | 6.9528 | 1 | 2 |
| R | F | 0.7170 | 6.7343 | 1 | 2 |
| R | S | 0.3061 | 6.1259 | 1 | 3 |
| R | H | 0.2083 | 6.5884 | 1 | 3 |
| R | N | 0.4170 | 6.3812 | 1 | 3 |
| R | P | 0.3204 | 6.3226 | 1 | 3 |
| R | C | 0.3125 | 6.2817 | 1 | 3 |

Table S4 continued from previous page

| Residue $i^{\text{th}}$ | Residue $j^{\text{th}}$ | $\epsilon$ (kcal mol$^{-1}$) | $\sigma$ (Å) | $\nu$ | $\mu$ |
| --- | --- | --- | --- | --- | --- |
| **R** | **I** | 0.2547 | 6.8804 | 1 | 3 |
| **S** | **S** | 0.1123 | 5.4127 | 1 | 4 |
| **S** | **H** | 0.3785 | 5.8752 | 1 | 3 |
| **S** | **N** | 0.2274 | 5.6680 | 1 | 3 |
| **S** | **P** | 0.1271 | 5.6094 | 1 | 3 |
| **S** | **C** | 0.1190 | 5.5685 | 1 | 3 |
| **S** | **I** | 0.0590 | 6.1672 | 1 | 4 |
| **T** | **T** | 0.0586 | 5.8891 | 1 | 4 |
| **T** | **R** | 0.2802 | 6.3641 | 1 | 3 |
| **T** | **A** | 0.0749 | 5.5796 | 1 | 4 |
| **T** | **D** | 0.1206 | 5.8563 | 1 | 3 |
| **T** | **E** | 0.1280 | 6.0334 | 1 | 3 |
| **T** | **Y** | 0.3654 | 6.3113 | 1 | 3 |
| **T** | **V** | 0.0366 | 6.0775 | 1 | 5 |
| **T** | **L** | 0.0414 | 6.2116 | 1 | 5 |
| **T** | **Q** | 0.2063 | 6.0835 | 1 | 3 |
| **T** | **W** | 0.4850 | 6.4778 | 1 | 3 |
| **T** | **F** | 0.3428 | 6.2593 | 1 | 3 |
| **T** | **S** | 0.0854 | 5.6509 | 1 | 4 |
| **T** | **H** | 0.3517 | 6.1134 | 1 | 3 |
| **T** | **N** | 0.2006 | 5.9062 | 1 | 3 |
| **T** | **P** | 0.1003 | 5.8476 | 1 | 4 |
| **T** | **C** | 0.0921 | 5.8067 | 1 | 4 |
| **T** | **I** | 0.0321 | 6.4054 | 1 | 5 |
| **V** | **V** | 0.0147 | 6.2660 | 1 | 6 |
| **V** | **L** | 0.0194 | 6.4000 | 1 | 6 |
| **V** | **Q** | 0.1844 | 6.2719 | 1 | 3 |
| **V** | **W** | 0.4642 | 6.6663 | 1 | 3 |

Table S4 continued from previous page

| Residue $i^{\text{th}}$ | Residue $j^{\text{th}}$ | $\epsilon$ (kcal mol$^{-1}$) | $\sigma$ (Å) | $\nu$ | $\mu$ |
| --- | --- | --- | --- | --- | --- |
| V | F | 0.3220 | 6.4478 | 1 | 3 |
| V | S | 0.0635 | 5.8393 | 1 | 4 |
| V | H | 0.3297 | 6.3019 | 1 | 3 |
| V | N | 0.1786 | 6.0947 | 1 | 3 |
| V | P | 0.0784 | 6.0361 | 1 | 4 |
| V | C | 0.0702 | 5.9952 | 1 | 4 |
| V | I | 0.0102 | 6.5938 | 1 | 7 |
| W | W | 0.8031 | 7.0666 | 1 | 2 |
| W | F | 0.7703 | 6.8481 | 1 | 2 |
| W | S | 0.5105 | 6.2396 | 1 | 3 |
| W | H | 0.7632 | 6.7022 | 1 | 2 |
| W | N | 0.6198 | 6.4950 | 1 | 2 |
| W | P | 0.5246 | 6.4364 | 1 | 3 |
| W | C | 0.5169 | 6.3955 | 1 | 3 |
| W | I | 0.4599 | 6.9941 | 1 | 3 |
| Y | Y | 0.6458 | 6.7336 | 1 | 2 |
| Y | V | 0.3447 | 6.4998 | 1 | 3 |
| Y | L | 0.3492 | 6.6339 | 1 | 3 |
| Y | Q | 0.5050 | 6.5057 | 1 | 3 |
| Y | W | 0.7929 | 6.9001 | 1 | 2 |
| Y | F | 0.6231 | 6.6816 | 1 | 2 |
| Y | S | 0.3908 | 6.0732 | 1 | 3 |
| Y | H | 0.6424 | 6.5357 | 1 | 2 |
| Y | N | 0.4996 | 6.3285 | 1 | 3 |
| Y | P | 0.4048 | 6.2699 | 1 | 3 |
| Y | C | 0.3971 | 6.2290 | 1 | 3 |
| Y | I | 0.3404 | 6.8277 | 1 | 3 |

##### SIII. STATISTICAL UNCERTAINTY QUANTIFICATION AND TRAJECTORY BLOCK-AVERAGING

This section details the mathematical framework used to evaluate statistical uncertainties, block-averaged standard errors, and confidence intervals across all molecular dynamics simulation trajectories.

###### A. Error Estimation for Correlated Molecular Dynamics Trajectories

In coarse-grained molecular dynamics simulations performed under Langevin dynamics, physical observables (such as intermolecular contact counts and pairwise interaction energies) are evaluated across discrete trajectory frames separated by time interval $\Delta t_{\text{sample}}$. Successive trajectory snapshots are correlated in time; consequently, the naive standard error of the mean underestimates the true statistical uncertainty if temporal correlation is neglected.

To obtain rigorous, unbiased standard errors for all trajectory-derived observables, we employ the trajectory block-averaging technique. For a stationary time series of $N$ sampled frames $\{x_1, x_2, \dots, x_N\}$ with sample mean $\bar{x} = \frac{1}{N} \sum_{k=1}^N x_k$, the trajectory is partitioned into $M$ non-overlapping blocks of length $B$ such that $N = M \cdot B$. The individual block averages are:

$$\bar{x}_b = \frac{1}{B} \sum_{k=(b-1)B+1}^{bB} x_k \quad (b = 1, \dots, M) \quad (\text{S10})$$

For sufficiently large block sizes $B$, the block averages become mutually independent and normally distributed by the Central Limit Theorem. The block-averaged standard error of the mean (SEM) as a function of block size $B$ is given by:

$$\text{SEM}(B) = \sqrt{\frac{1}{M(M-1)} \sum_{b=1}^M (\bar{x}_b - \bar{x})^2} \quad (\text{S11})$$

We evaluate block sizes across $B \in [2, 5, 10, 20, 25, 50, 100]$ snapshots and determine the asymptotic plateau value $\text{SEM} = \max_B \text{SEM}(B)$, ensuring that all reported standard errors rigorously account for molecular relaxation times. The 95% confidence interval (CI) is evaluated as $\bar{x} \pm 1.96 \times \text{SEM}$.

#### B. Ensemble and Multi-Replicate Variance

For massive macromolecular assemblies (such as the 2521-residue PIEZO1 monomer in Figure 4), independent production trajectories were conducted across separate initial configurations and random velocity seeds (replicate runs *rep1*, *rep2*, and *platform*). For any observable $y$ (e.g., the 50-residue sliding-window interaction energy), the ensemble replicate mean $\bar{y}$ and sample standard deviation $s_{\text{rep}}$ across $K$ replicates ($K \in [2, 3]$) are evaluated as:

$$\bar{y} = \frac{1}{K} \sum_{r=1}^K y_r, \quad s_{\text{rep}} = \sqrt{\frac{1}{K-1} \sum_{r=1}^K (y_r - \bar{y})^2}, \quad \text{SEM}_{\text{rep}} = \frac{s_{\text{rep}}}{\sqrt{K}} \quad (\text{S12})$$

#### C. Uncertainty Propagation for Specificity Ratios

To evaluate binding specificity (Figures 3 and 4), the specificity ratio $S$ of on-target interaction energy $E_{\text{on}}$ relative to off-target cross-reactivity $E_{\text{off}}$ is defined as:

$$S = \frac{|E_{\text{on}}|}{|E_{\text{off}}|} \quad (\text{S13})$$

Assuming independent errors in the on-target and off-target simulation ensembles, the propagated standard error $\sigma_S$ is determined via first-order Taylor expansion:

$$\sigma_S = S \sqrt{\left( \frac{\sigma_{\text{on}}}{|E_{\text{on}}|} \right)^2 + \left( \frac{\sigma_{\text{off}}}{|E_{\text{off}}|} \right)^2} \quad (\text{S14})$$

#### D. Critical Scaling Regression Covariance for CHERP Condensates

Direct coexistence slab simulations across temperatures $T \in [300, 390 \text{ K}]$ yield coexistence densities $\rho_{\text{dilute}}(T)$ and $\rho_{\text{dense}}(T)$ (Figure 5). The critical solution temperature $T_c$ and critical density $\rho_c$ are determined by non-linear least-squares fitting using the Levenberg–Marquardt algorithm to the universal 3D Ising scaling law and the law of rectilinear diameters:

$$\rho_{\text{dense}}(T) - \rho_{\text{dilute}}(T) = 2B \left( 1 - \frac{T}{T_c} \right)^\beta \quad (\text{S15})$$

$$\frac{\rho_{\text{dense}}(T) + \rho_{\text{dilute}}(T)}{2} = \rho_c + A (T_c - T) \quad (\text{S16})$$

where $\beta = 0.326$ is the exact 3D Ising critical exponent. The asymptotic standard errors for the fit parameters $(\sigma_{T_c}, \sigma_B, \sigma_{\rho_c}, \sigma_A)$ are extracted from the diagonal elements of the parameter covariance matrix $\mathbf{V}$:

$$\sigma_{p_i} = \sqrt{V_{ii}}, \quad \mathbf{V} = (\mathbf{J}^T \mathbf{W} \mathbf{J})^{-1} \quad (\text{S17})$$

where $\mathbf{J}$ is the Jacobian matrix of partial derivatives and $\mathbf{W}$ is the diagonal weight matrix corresponding to inverse simulation variances. The uncertainty in the critical temperature shift $\Delta T_c = T_c^{\text{pep}} - T_c^{\text{pure}}$ is propagated as:

$$\sigma_{\Delta T_c} = \sqrt{\sigma_{T_c, \text{pep}}^2 + \sigma_{T_c, \text{pure}}^2} \quad (\text{S18})$$

#### SIV. COMPARATIVE EVALUATION STATISTICS, SUPPLEMENTARY TABLES, AND SUPPLEMENTARY FIGURES

This section compiles the comprehensive numerical evaluation tables (Tables S5–S13) and high-resolution supplementary figures (Figs. S1, S2, and S3) providing statistical verification for the designs, evolutionary trajectories, and multi-scale simulations reported in Figures 2 through 5 of the main manuscript.

##### A. Machine-Learning Comparisons and Directed Evolution Runtime (Figure 2)

In directed evolution campaigns, each generation evaluates a population of 15 candidate peptides. For each candidate peptide, two independent MD simulations are executed: (1) a target-binding and off-target cross-reactivity simulation ($f_{\text{bind}}$ and $f_{\text{cross}}$), and (2) a multi-copy peptide-only self-aggregation simulation in solvent ($f_{\text{agg}}$), totaling 30 distinct physical simulations per generation (15 candidates × 2 simulation types). Across a 50-round campaign, this accumulates to a total of 1,500 independent simulation trajectories (750 binding and 750 self-aggregation runs, 40 ns per trajectory). Each individual simulation was allocated 16 MPI tasks (16 CPU cores per simulation job). When submitted concurrently across cluster nodes in SLURM (15 × 2 × 16 = 480 total parallel MPI tasks per generation), each generation completed in approximately 19 minutes of wall-clock time, yielding an overall campaign turnaround of ∼ 16 hours. Users can flexibly adjust the number of MPI tasks per simulation or candidate population size to scale execution throughput with available cluster resources. For comparison, sequence-only language models like *pepMLM* execute inference in seconds on a single GPU, whereas structural diffusion pipelines (*RFdiffusion* combined with ProteinMPNN and AlphaFold2 filtering) typically require ∼ 15–40 minutes per batch of validated designs on GPU hardware. PepSpace achieves comparable per-batch turnaround on accessible CPU infrastructure while providing explicit thermodynamic ensemble sampling and multi-objective negative-selection filtering. Crucially, PepSpace explores regions of sequence space that generative models do not reach under this physical objective, actively optimizing intermolecular interactions while filtering out self-aggregation. Table S5 and Supplementary Figure S3 compile the physical interaction energies per peptide residue ($|\bar{E}_{\text{inter}}|$, kcal/mol per amino acid) and trajectory block-averaged standard errors (SEM) across all 16 generative benchmark candidates (4 *pepMLM* and 4 *RFdiffusion* designs per receptor) on AMHR2 and NCAM1, highlighting the primary experimental leads (*pepMLM-1* and *pepMLM-2*).

Supplementary Figure S1 displays the 50-round evolutionary trajectory convergence envelopes alongside sequence diversity preservation (mean pairwise Hamming distance) across both targets. Table S6 and Supplementary Figure S2 document the binding interaction energy enhancements of the top-5 evolved PepSpace candidates compared to the baseline *pepMLM* designs, confirming 16.2-fold (AMHR2) and 179.6-fold (NCAM1) enhancements ($p < 10^{-15}$).

TABLE S5. **Statistical metrics for generative machine-learning benchmarks (*pepMLM* and *RFdiffusion*) on AMHR2 and NCAM1 (Table S5, complementing Figure 2a,b and Supplementary Figure S3).** Tabulated are the target receptor, candidate peptide designation, absolute coarse-grained MD interaction energy per residue ($|\bar{E}_{\text{inter}}|$, kcal/mol/AA), and trajectory block-averaged standard error (SEM). Primary experimental benchmark leads are highlighted in parentheses.

| Target Receptor | Peptide Designation | $\vert \bar{E}_{\text{inter}}\vert $ (kcal/mol/AA) | SEM |
| --- | --- | --- | --- |
| AMHR2 | AMHR2-pepMLM-01 | 0.0028 | 0.0001 |
| AMHR2 | AMHR2-pepMLM-02 | 0.0097 | 0.0004 |
| AMHR2 | AMHR2-pepMLM-03 (*pepMLM-1*) | 0.0073 | 0.0003 |
| AMHR2 | AMHR2-pepMLM-04 | 0.0067 | 0.0003 |
| AMHR2 | AMHR2-RFD-01 (*RFD-1*) | 0.0005 | 0.00003 |
| AMHR2 | AMHR2-RFD-02 | 0.0006 | 0.00003 |
| AMHR2 | AMHR2-RFD-03 | 0.0037 | 0.0002 |
| AMHR2 | AMHR2-RFD-04 | 0.0025 | 0.0001 |
| NCAM1 | NCAM1-pepMLM-01 | 0.0007 | 0.00004 |
| NCAM1 | NCAM1-pepMLM-02 (*pepMLM-2*) | 0.0018 | 0.0001 |
| NCAM1 | NCAM1-pepMLM-03 | 0.0007 | 0.00004 |
| NCAM1 | NCAM1-pepMLM-04 | 0.0015 | 0.0001 |
| NCAM1 | NCAM1-RFD-01 | 0.0019 | 0.0001 |
| NCAM1 | NCAM1-RFD-02 (*RFD-2*) | 0.0017 | 0.0001 |
| NCAM1 | NCAM1-RFD-03 | 0.0019 | 0.0001 |
| NCAM1 | NCAM1-RFD-04 | 0.0016 | 0.0001 |

TABLE S6. **Statistical evaluation of top-5 evolved peptides versus benchmark *pepMLM* designs on AMHR2 and NCAM1 (Table S6, complementing Figure 2e,f).** Tabulated are the target receptor, candidate peptide designation, absolute physical binding energy per residue ($|\bar{E}_{\text{inter}}|$, kcal/mol/AA), and trajectory block-averaged standard error (SEM), and fold improvement over baseline. All evolved improvements are statistically significant by Welch’s $t$-test ($p < 10^{-15}$).

| Target Receptor | Peptide Designation | $\vert \bar{E}_{\text{inter}}\vert $ (kcal/mol/AA) | SEM | Fold Gain |
| --- | --- | --- | --- | --- |
| AMHR2 | AMHR2-pepMLM-03 (*pepMLM-1*, Baseline) | 0.0073 | 0.0003 | 1.0× (Reference) |
| AMHR2 | top-01 AMHR2 | 0.118 | 0.001 | 16.2× |
| AMHR2 | top-02 AMHR2 | 0.115 | 0.002 | 15.8× |
| AMHR2 | top-03 AMHR2 | 0.114 | 0.002 | 15.6× |
| AMHR2 | top-04 AMHR2 | 0.110 | 0.001 | 15.0× |
| AMHR2 | top-05 AMHR2 | 0.107 | 0.002 | 14.7× |
| NCAM1 | NCAM1-pepMLM-02 (*pepMLM-2*, Baseline) | 0.0018 | 0.0001 | 1.0× (Reference) |
| NCAM1 | top-01 NCAM1 | 0.325 | 0.014 | 179.6× |
| NCAM1 | top-02 NCAM1 | 0.289 | 0.012 | 160.0× |
| NCAM1 | top-03 NCAM1 | 0.274 | 0.010 | 151.5× |
| NCAM1 | top-04 NCAM1 | 0.262 | 0.010 | 144.6× |
| NCAM1 | top-05 NCAM1 | 0.249 | 0.009 | 137.5× |

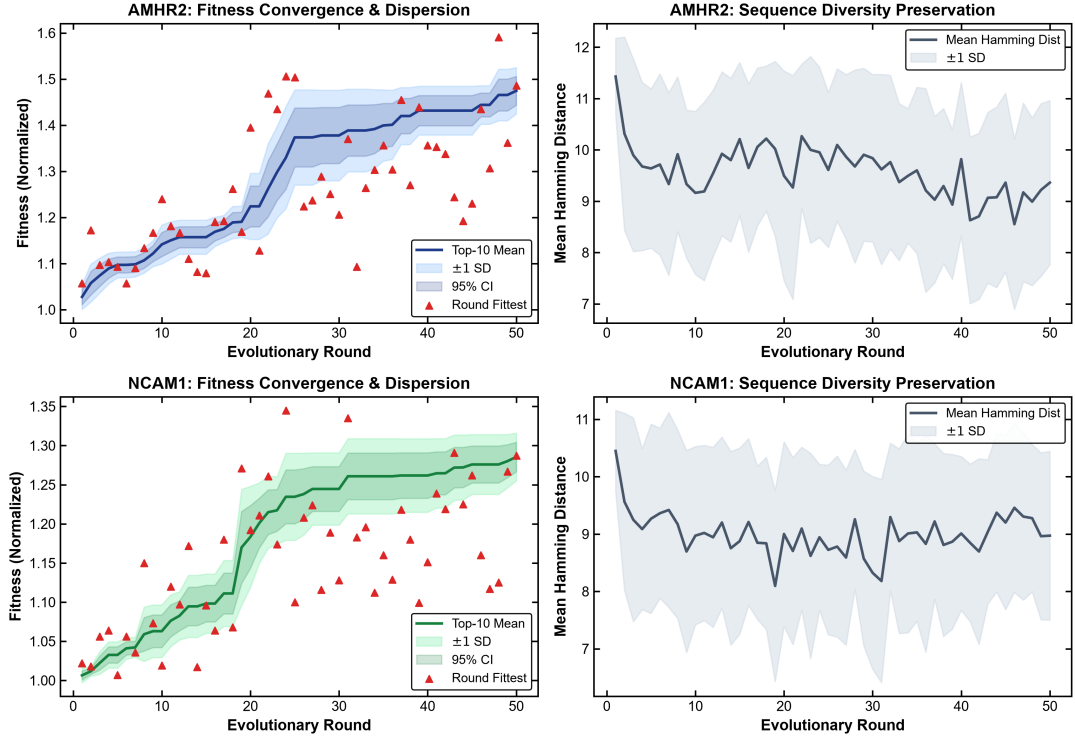

FIG. S1. **Evolutionary trajectory convergence, error envelopes, and population diversity preservation (Figure S1, complementing Figure 2c,d).** **Left panels:** Mean composite fitness of the top-10 candidate population across 50 evolutionary rounds for AMHR2 (top) and NCAM1 (bottom); solid curve indicates the top-10 mean, shaded lighter band shows $\pm 1$ standard deviation ($s$), shaded darker band indicates the 95% confidence interval, and red triangles indicate the fittest candidate in each generation. **Right panels:** Population sequence diversity quantified by the mean pairwise Hamming distance across the 15-candidate generation pool (solid grey line) with $\pm 1$ standard deviation envelope (light grey band).

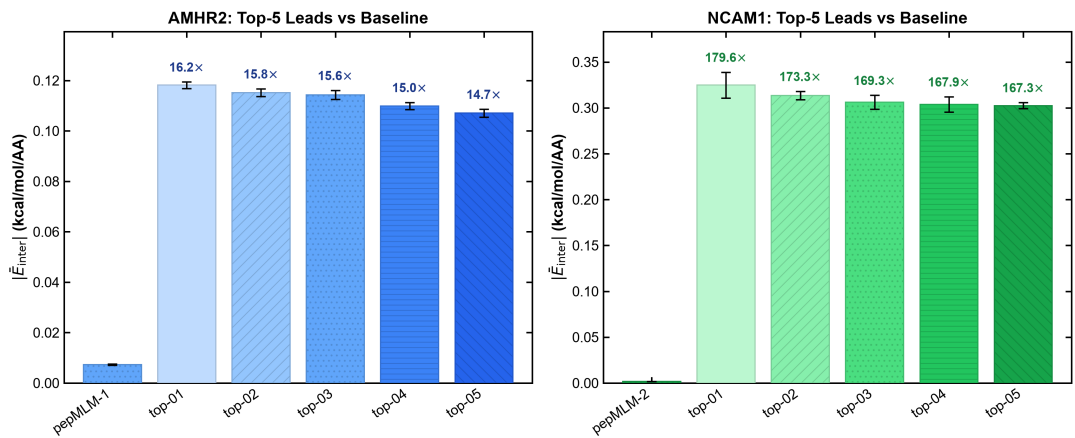

FIG. S2. **Linear-scale binding interaction energy comparison with explicit error bars for top-5 evolved peptides versus pepMLM baselines (Figure S2, complementing Figure 2e,f).** Bar charts showing absolute interaction energies per residue ($|\bar{E}_{\text{inter}}|$, kcal/mol/AA) on AMHR2 (left) and NCAM1 (right); error bars represent block-averaged standard errors (SEM). Baseline experimental *pepMLM* designs are shown with respective hatch and color matching main text Figure 2; colored hatched bars indicate the top-5 evolved PepSpace candidates, and text annotations indicate fold affinity enhancements (16.2× on AMHR2, 179.6× on NCAM1; Welch’s *t*-test $p < 10^{-15}$).

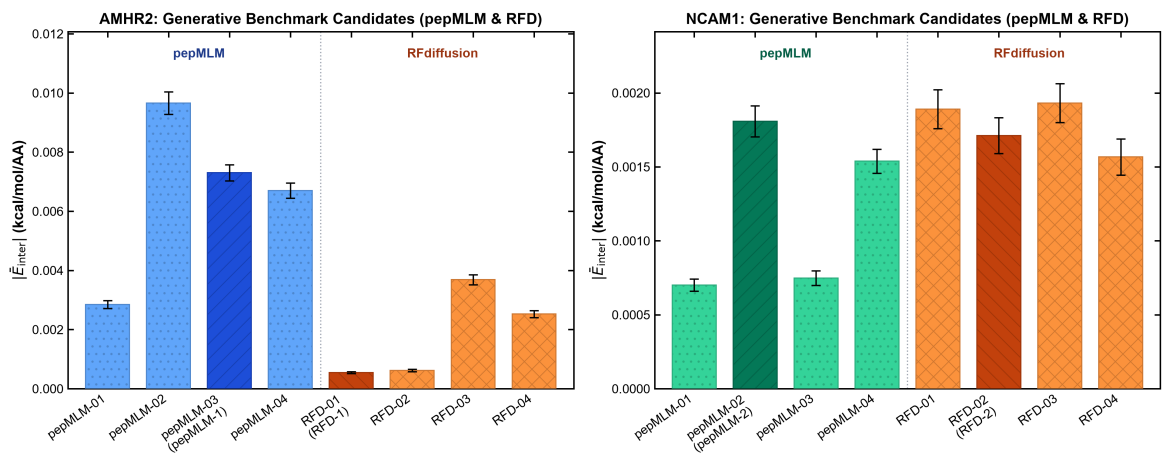

FIG. S3. **Linear-scale binding interaction energy comparison across all 16 generative benchmark candidates on AMHR2 and NCAM1 (Figure S3, complementing Figure 2a,b and Table S5).** Absolute coarse-grained MD interaction energies per residue ($|\bar{E}_{\text{inter}}|$, kcal/mol/AA) for 8 candidates per receptor (4 designed with *pepMLM* and 4 designed with *RFdiffusion*); error bars represent block-averaged standard errors of the mean (SEM). Colors and hatching correspond to the unified design schema; primary experimental benchmark leads (*pepMLM*-1: AMHR2-pepMLM-03, and *pepMLM*-2: NCAM1-pepMLM-02) are highlighted.

##### B. Cross-Reactivity and Target Specificity (Figure 3)

Table S7 reports quantitative on-target ($|E_{\text{on}}|$) versus off-target ($|E_{\text{off}}|$) cross-reactivity metrics. Table S8 summarizes the domain-level partitioning of physical binding energies across the structural domains of AMHR2 and NCAM1, demonstrating that high-affinity binding is concentrated at specific structured domains while non-specific background binding is suppressed.

TABLE S7. **Quantitative on-target versus off-target cross-reactivity metrics (Table S7, complementing Figure 3a,b).** Tabulated are the candidate peptide designation, cognate target receptor, on-target binding energy per residue ($|E_{\text{on}}|$, kcal/mol/AA), and off-target cross-reactive binding energy ($|E_{\text{off}}|$, kcal/mol/AA). All reported energies denote mean ± SEM; off-target receptor corresponds to NCAM1 for AMHR2 candidates and AMHR2 for NCAM1 candidates.

| Candidate Peptide | Cognate Target | $\vert E_{\text{on}}\vert $ (kcal/mol/AA) | $\vert E_{\text{off}}\vert $ (kcal/mol/AA) |
| --- | --- | --- | --- |
| top-01 AMHR2 | AMHR2 | $0.118 \pm 0.001$ | $0.271 \pm 0.007$ |
| AMHR2-pepMLM-03 (*pepMLM-1*) | AMHR2 | $0.0073 \pm 0.0003$ | $0.009 \pm 0.0004$ |
| top-01 NCAM1 | NCAM1 | $0.325 \pm 0.014$ | $0.112 \pm 0.002$ |
| NCAM1-pepMLM-02 (*pepMLM-2*) | NCAM1 | $0.0018 \pm 0.0001$ | $0.001 \pm 0.0001$ |

TABLE S8. **Domain-resolved interaction energy intensity for cognate and off-target peptides across AMHR2 and NCAM1 structural domains (Table S8, complementing Figure 3c,d).** Tabulated are the target receptor, candidate peptide, structural domain with residue boundaries, and mean contact interaction energy per target residue (mean ± trajectory SEM, kcal/mol/AA).

| Target Receptor | Candidate Peptide | Structural Domain | Mean Energy (kcal/mol/AA) |
| --- | --- | --- | --- |
| AMHR2 | top-01 AMHR2 | N-term Loop (1–35) | $0.107 \pm 0.001$ |
| AMHR2 | top-01 AMHR2 | Core β-Sheet (36–95) | $0.068 \pm 0.001$ |
| AMHR2 | top-01 AMHR2 | C-term Region (96–128) | $0.129 \pm 0.001$ |
| AMHR2 | top-01 NCAM1 | N-term Loop (1–35) | $0.099 \pm 0.002$ |
| AMHR2 | top-01 NCAM1 | Core β-Sheet (36–95) | $0.044 \pm 0.001$ |
| AMHR2 | top-01 NCAM1 | C-term Region (96–128) | $0.170 \pm 0.003$ |
| NCAM1 | top-01 AMHR2 | Ig1 Domain (1–96) | $0.011 \pm 0.0003$ |
| NCAM1 | top-01 AMHR2 | Ig2 Domain (97–190) | $0.003 \pm 0.0001$ |
| NCAM1 | top-01 AMHR2 | Ig3 Domain (191–285) | $0.025 \pm 0.0006$ |
| NCAM1 | top-01 AMHR2 | Ig4 Domain (286–380) | $0.139 \pm 0.0034$ |
| NCAM1 | top-01 AMHR2 | Ig5 Domain (381–490) | $0.025 \pm 0.0006$ |
| NCAM1 | top-01 AMHR2 | FnIII Domains (491–709) | $0.013 \pm 0.0003$ |
| NCAM1 | top-01 NCAM1 | Ig1 Domain (1–96) | $0.066 \pm 0.0029$ |
| NCAM1 | top-01 NCAM1 | Ig2 Domain (97–190) | $0.002 \pm 0.0001$ |
| NCAM1 | top-01 NCAM1 | Ig3 Domain (191–285) | $0.102 \pm 0.0044$ |
| NCAM1 | top-01 NCAM1 | Ig4 Domain (286–380) | $0.030 \pm 0.0013$ |
| NCAM1 | top-01 NCAM1 | Ig5 Domain (381–490) | $0.090 \pm 0.0039$ |
| NCAM1 | top-01 NCAM1 | FnIII Domains (491–709) | $0.015 \pm 0.0007$ |

##### C. PIEZO1 Beam Targeting and Replicate Convergence (Figure 4)

Table S9 summarizes multi-replicate binding statistics across independent trajectories totaling 20.0 μs of aggregate sampling, confirming a 41.4-fold (PIEZO1-01) and 46.4-fold (PIEZO1-02) signal-to-noise specificity enrichment for the designated intracellular beam epitope (Window 28, residues 1373–1402) over the remaining 2500-residue channel body. Table S10 confirms that cross-reactivity on non-cognate receptors AMHR2 and NCAM1 is attenuated by up to 22-fold.

TABLE S9. **Whole-protein multi-replicate interaction energy statistics for PIEZO1 designed peptides (Table S9, complementing Figure 4c,d).** Tabulated are the candidate peptide, target epitope interaction energy (Window 28, mean ± SEM, kcal/mol across $K = 3$ replicates for PIEZO1-01 and $K = 2$ for PIEZO1-02), non-target background window energy (mean ± SD across all 50 non-target windows), and the resulting on-target specificity enrichment ratio.

| Candidate Peptide | Target Energy (kcal/mol) | Off-Target Energy (kcal/mol) | Enrichment Ratio |
| --- | --- | --- | --- |
| PIEZO1-01 | $-1.316 \pm 0.165$ | $-0.032 \pm 0.025$ | 41.4× |
| PIEZO1-02 | $-1.602 \pm 0.130$ | $-0.035 \pm 0.027$ | 46.4× |

TABLE S10. **Off-target cross-reactivity statistics of PIEZO1 designed peptides on non-cognate receptors AMHR2 and NCAM1 (Table S10, complementing Figure 4g).** Tabulated are the tested receptor, candidate peptide designation, binding status, absolute interaction energy per residue ($|\bar{E}_{\text{inter}}|$, mean ± SEM, kcal/mol/AA), and off-target attenuation fold-factor relative to the cognate fitted binder.

| Receptor | Candidate Peptide | Status | $\vert \bar{E}_{\text{inter}}\vert $ (kcal/mol/AA) | Attenuation |
| --- | --- | --- | --- | --- |
| AMHR2 | top-01 AMHR2 | Cognate | $0.118 \pm 0.001$ | 1.0× (Reference) |
| AMHR2 | PIEZO1-01 | Off-target | $0.009 \pm 0.0003$ | 12.7× Reduction |
| AMHR2 | PIEZO1-02 | Off-target | $0.010 \pm 0.0004$ | 11.6× Reduction |
| NCAM1 | top-01 NCAM1 | Cognate | $0.325 \pm 0.014$ | 1.0× (Reference) |
| NCAM1 | PIEZO1-01 | Off-target | $0.074 \pm 0.003$ | 4.4× Reduction |
| NCAM1 | PIEZO1-02 | Off-target | $0.015 \pm 0.001$ | 22.4× Reduction |

###### D. CHERP Phase Separation Modulators (Figure 5)

Table S11 compiles the critical point parameters and 3D Ising scaling regression diagnostics from direct coexistence slab simulations across $T \in [300, 390 \text{ K}]$, yielding critical solution temperatures of $T_c = 390.5 \text{ K}$ for pure CHERP, $381.2 \text{ K}$ for CHERP-02 ($\Delta T_c = -9.3 \text{ K}$), and $377.2 \text{ K}$ for CHERP-01 ($\Delta T_c = -13.3 \text{ K}$). Table S12 details the homotypic CHERP–CHERP interaction energy per pair ($\bar{E}_{C-C}^{\text{pair}}$, kcal/mol per pair) at $T = 370 \text{ K}$, evaluated globally across all 300 intermolecular pairs and strictly within the 476–576 condensation hot spot, demonstrating 21.1% global and 22.4% hotspot energy attenuation by CHERP-01. Summing the per-residue intermolecular interaction energies $\sum_{i=476}^{576} E_{C-C}(i)$ yields exactly the $-0.671 \text{ kcal/mol per pair}$ reported in Table S12, corresponding to the integrated area under the 1D profile displayed in Figure 5d. Table S13 reports cross-reactivity suppression on non-cognate receptors AMHR2 and NCAM1.

TABLE S11. **Critical point parameters and 3D Ising scaling regression diagnostics for CHERP phase diagrams (Table S11, complementing Figure 5b).** Tabulated are the system condition, critical temperature ($T_c$, K), critical density ($\rho_c$, mg/mL), Ising width coefficient ($B$), rectilinear slope ($A$), and critical temperature shift ($\Delta T_c$, K) relative to pure CHERP from direct coexistence simulations.

| System | $T_c$ (K) | $\rho_c$ (mg/mL) | Ising $B$ | Slope $A$ | $\Delta T_c$ (K) |
| --- | --- | --- | --- | --- | --- |
| Pure CHERP | 390.5 | 80.9 | 217.7 | 0.618 | 0.0 (Reference) |
| CHERP-02 | 381.2 | 77.2 | 239.8 | 0.959 | −9.3 |
| CHERP-01 | 377.2 | 78.7 | 256.3 | 1.098 | −13.3 |

TABLE S12. **Homotypic CHERP–CHERP intermolecular interaction energy per pair at 370 K (Table S12, complementing Figure 5f,g).** Tabulated are the system condition, total homotypic interaction energy per pair ($|\bar{E}_{\text{inter}}|$, mean ± SEM, kcal/mol per pair), percentage total reduction, condensation hotspot (residues 476–576) per-pair energy (mean ± SEM), and hotspot percentage reduction across the 300 intermolecular pairs.

| System | Total Energy (kcal/mol/pair) | Total $\Delta$ | Hotspot Energy (kcal/mol/pair) | Hotspot $\Delta$ |
| --- | --- | --- | --- | --- |
| Pure CHERP | $2.670 \pm 0.053$ | 0.0% (Reference) | $0.671 \pm 0.013$ | 0.0% (Reference) |
| CHERP-01 | $2.107 \pm 0.092$ | −21.1% | $0.521 \pm 0.023$ | −22.4% |
| CHERP-02 | $2.272 \pm 0.033$ | −14.9% | $0.574 \pm 0.008$ | −14.5% |

TABLE S13. **Off-target cross-reactivity statistics of CHERP designed peptides on non-cognate receptors AMHR2 and NCAM1 (Table S13, complementing Figure 5h).** Tabulated are the tested receptor, candidate peptide designation, binding status, absolute interaction energy per residue ($|\bar{E}_{\text{inter}}|$, mean ± SEM, kcal/mol/AA), and off-target attenuation relative to the cognate fitted binder.

| Receptor | Candidate Peptide | Status | $\vert \bar{E}_{\text{inter}}\vert $ (kcal/mol/AA) | Attenuation |
| --- | --- | --- | --- | --- |
| AMHR2 | top-01 AMHR2 | Cognate | $0.118 \pm 0.001$ | 1.0× (Reference) |
| AMHR2 | CHERP-01 | Off-target | $0.009 \pm 0.0003$ | 13.9× Reduction |
| AMHR2 | CHERP-02 | Off-target | $0.011 \pm 0.0004$ | 10.5× Reduction |
| NCAM1 | top-01 NCAM1 | Cognate | $0.325 \pm 0.014$ | 1.0× (Reference) |
| NCAM1 | CHERP-01 | Off-target | $0.038 \pm 0.001$ | 8.5× Reduction |
| NCAM1 | CHERP-02 | Off-target | $0.015 \pm 0.001$ | 22.4× Reduction |

- [1] Tejedor, A. R. *et al.* Chemically informed coarse-graining of electrostatic forces in charge-rich biomolecular condensates. *ACS Central Science* **11**, 302–321 (2025).
- [2] Wang, X., Ramírez-Hinestrosa, S., Dobnikar, J. & Frenkel, D. The Lennard-Jones potential: when (not) to use it. *Physical Chemistry Chemical Physics* **22**, 10624–10633 (2020).
